# Magnitude and kinetics of T cell and antibody responses during H1N1pdm09 infection in outbred and inbred Babraham pigs

**DOI:** 10.1101/2020.10.28.346767

**Authors:** Matthew Edmans, Adam McNee, Emily Porter, Eleni Vatzia, Basu Paudyal, Veronica Martini, Simon Gubbins, Ore Francis, Ross Harley, Amy Thomas, Rachel Burt, Sophie Morgan, Anna Fuller, Andrew Sewell, sLoLA Influenza Dynamics consortium, Bryan Charleston, Mick Bailey, Elma Tchilian

**Affiliations:** The Pirbright Institute, Pirbright, UK; Bristol Veterinary School, University of Bristol, Langford, UK; Division of Infection and Immunity, Cardiff University School of Medicine, Cardiff, UK

**Author notes:** The sLoLa Influenza Dynamics consortium is (in alphabetical order): Mario Aramouni, Mick Bailey, Amy Boyd, Sharon Brookes, Ian Brown, Becky Clark, Bryan Charleston, Catherine Charreyre, Margo Chase-Topping, Federica Di-Palma, Matthew Edmans, Graham Etherington, Helen Everett, Ore Francis, Simon Frost, Sarah Gilbert, Ross Harley, Barbara Holzer, Adam McNee, Angela Man, Veronica Martini, Sophie Morgan, Emily Porter, Jin Qi Fu, Amy Thomas, Elma Tchilian, Laurence Tiley, Pauline van Diemen, James Wood, Fei Xiang. Corresponding author Elma Tchilian. These authors contributed equally (Matthew Edmans, Adam McNee, Emily Porter, Bryan Charleston, Mick Bailey, Elma Tchilian).

## Abstract

We have used the pig, a large natural host animal for influenza with many physiological similarities to humans, to characterize αβ, γδ T cell and antibody (Ab) immune responses to the 2009 pandemic H1N1 virus infection. We evaluated the kinetic of virus infection and associated response in inbred Babraham pigs with identical MHC (Swine Leucocyte Antigen) and compared them to commercial outbred animals. High level of nasal virus shedding continued up to day 4-5 post infection followed by a steep decline and clearance of virus by day 9. Adaptive T cell and Ab responses were detectable from day 5-6 post infection reaching a peak at 9-14 days. γδ cells produced cytokines *ex vivo* at day 2 post infection, while virus specific IFNγ producing γδ T cells were detected from day 7 post infection. Analysis of NP tetramer specific and virus specific CD8 and CD4 T cells in blood, lung, lung draining lymph nodes and broncho-alveolar lavage (BAL) showed clear differences in cytokine production between these tissues. BAL contained the most highly activated CD8, CD4 and γδ cells producing large amounts of cytokines, which likely contribute to elimination of virus. The weak response in blood did not reflect the powerful local lung immune responses. The immune response in the Babraham pig following H1N1pdm09 influenza infection was comparable to that of outbred animals. The ability to utilize these two swine models together will provide unparalleled power to analyse immune responses to influenza.

## Introduction

Influenza viruses are a global health threat to humans and pigs, causing considerable morbidity and mortality. Frequent zoonotic crossover between pigs and humans contributes to the evolution of influenza viruses and can be a source for novel pandemic strains (Nelson and Vincent, 2015; Kaplan et al., 2020; Sun et al., 2020). The emergence of the 2009 pandemic H1N1 (H1N1pdm09) virus, which is now globally endemic in both pigs and humans, illustrates the importance of pigs in new outbreaks in humans (Smith et al., 2009). Influenza A virus (IAV) infection in pigs causes significant economic loss due to reduced weight gain, suboptimal reproductive performance and secondary infections. Immunization with inactivated influenza virus is currently the most effective way of inducing strain-specific neutralizing antibodies, directed against the surface glycoprotein haemagglutinin (HA). Because of the constant evolution of the virus, broadly cross-protective vaccines would be highly desirable and central to the control of influenza in both pigs and humans.

Animal models are essential to develop better vaccines and control strategies and to provide insight into human disease. Most models have limitations in recapitulating the full range of disease observed in humans. Mice, guinea pigs and non-human primates are not generally susceptible to natural routes of influenza infection and may require adapted strains, physiologic stressors and/or unnatural inoculation procedures (Bouvier and Lowen, 2010; Margine and Krammer, 2014; Hemmink et al., 2018; Mifsud et al., 2018). In contrast, pigs are an important, natural, large animal host for IAV and are infected by the same subtypes of H1N1 and H3N2 viruses as humans (Watson et al., 2015; Lewis et al., 2016). Pigs have a longer life span, are genetically, immunologically, physiologically and anatomically more like humans than small laboratory animals and have a comparable distribution of sialic acid receptors in the respiratory tract (Janke, 2014; Rajao and Vincent, 2015). Pigs exhibit similar clinical manifestations and pathogenesis when infected with IAV making them an excellent model to study immunity to influenza. Furthermore, we have defined the dynamics of H1N1pdm09 influenza virus transmission in pigs and demonstrated the utility of the pig model to test therapeutic antibody delivery platforms and vaccines (Canini et al., 2020; McNee et al., 2020).

Several inbred miniature pig breeds have been developed, including NIH and Yucatan, with defined swine leukocyte antigens (SLA type, the swine major histocompatibility complex) (Sachs et al., 1976; Choi et al., 2016). However, the inbred Babraham is the only example of a full-size inbred strain of pig, closely related to commercial breeds, making them an appropriate model to study diseases important to commercial pig production (Signer et al., 1999; Schwartz et al., 2018). The sharing of IAV strains between pigs and humans makes it an obvious species in which to study immunity to influenza and to test vaccines or therapeutic strategies prior to human clinical trials. In addition we have developed a toolset to study immune responses in Babrahams, including adoptive cell transfer and peptide SLA tetramers allowing us to study the fine specificity of immune responses (Binns et al., 1981; Tungatt et al., 2018).

Despite extensive knowledge of the role of T cells in protection against IAV in mice and humans, few studies in pigs have evaluated this in depth. The duration and magnitude of T cell and humoral responses has been assessed after swine H1N1, H1N2 and H3N2 challenges in pigs (Heinen et al., 2000; Larsen et al., 2000; Khatri et al., 2010; Talker et al., 2015; Talker et al., 2016). The frequency and activation status of leucocytes in local and systemic tissues was also determined after H1N1pdm09 infection (Schwaiger et al., 2019). However no detailed analysis of T cell immune responses in broncho-alveolar lavage (BAL) have been performed, a location which we have shown to contain tissue resident memory cells that are essential for heterosubtypic protection (Holzer et al., 2018a). Neither has there been a detailed analysis of T cell and antibody (Ab) immune responses to H1N1pdm09, although this continues to cross the species barrier from humans to pigs. H1N1pdm09 circulating in swine herds maintains antigenic similarity to human seasonal strains, providing a unique opportunity to use a virus affecting both humans and swine to examine immune responses induced by infection.

Here we characterized αβ, γδ T cell and Ab immune responses to H1N1pdm09 in local lung and systemic tissues in Babraham pigs and compared them to commercial outbred animals. These two pig models together will allow fine grain dissection of immune responses to IAV in a species which is a natural host for the virus and similar in many respects to humans.

## Materials and methods

### Animals and influenza H1N1pdm09 challenge

The animal experiments were approved by the ethical review processes at the Pirbright Institute and Bristol and conducted according to the UK Government Animal (Scientific Procedures) Act 1986 under project licence P47CE0FF2. Both Institutes conform to the ARRIVE guidelines.

Thirty two outbred old Landrace x Hampshire cross (from a commercial high health status herd) and 56 inbred Babraham pigs (bred at Animal Plant Health Agency, APHA Weybridge, UK) were screened for absence of influenza A infection by matrix gene real time RT-PCR and for antibody-free status by HAI using four swine influenza virus antigens - H1N1pdm09, H1N2, H3N2 and avian-like H1N1. The average age of the outbred pigs 7 days before the challenge was 8.7 weeks and of the Babrahams 8.3 weeks. Pigs were challenged intra-nasally with 1 × 10^7^ PFU of MDCK grown swine A(H1N1)pdm09 isolate, A/swine/England/1353/2009, derived from the 2009 pandemic virus, swine clade 1A.3. (H1N1pdm09) in a total of 4 ml (2 ml per nostril) using a mucosal atomisation device MAD300 (MAD, Wolfe-Tory Medical). Two experiments with outbred (OB) pigs (referred to as OB1 and OB2) and two with inbred Babraham (BM) pigs (referred to as BM1 and BM2) were performed (**Fig. 1A**). In each experiment one pig was culled on days 1 to 7, 9, 11 and 13 post infection and a *post-mortem* examination performed with collection of tissue samples. Uninfected controls were sampled: two on the day prior to infection and two at day 8 post infection. Two naïve pigs (referred to as in-contact) were co-housed with the directly challenged pigs in experiments OB1, OB2, BM1, BM2 and culled at days 11 and 13 post infection together with the last two directly challenged pigs. A fifth experiment was performed with Babraham pigs (experiment BM3) in which 3 were culled on days 6, 7, 13, 14, 20 and 21 post infection (**Fig. 1A**). In the BM3 experiment 6 control animals were included, 3 of which were culled one day before and 3 on the day of infection.

**Figure 1.**
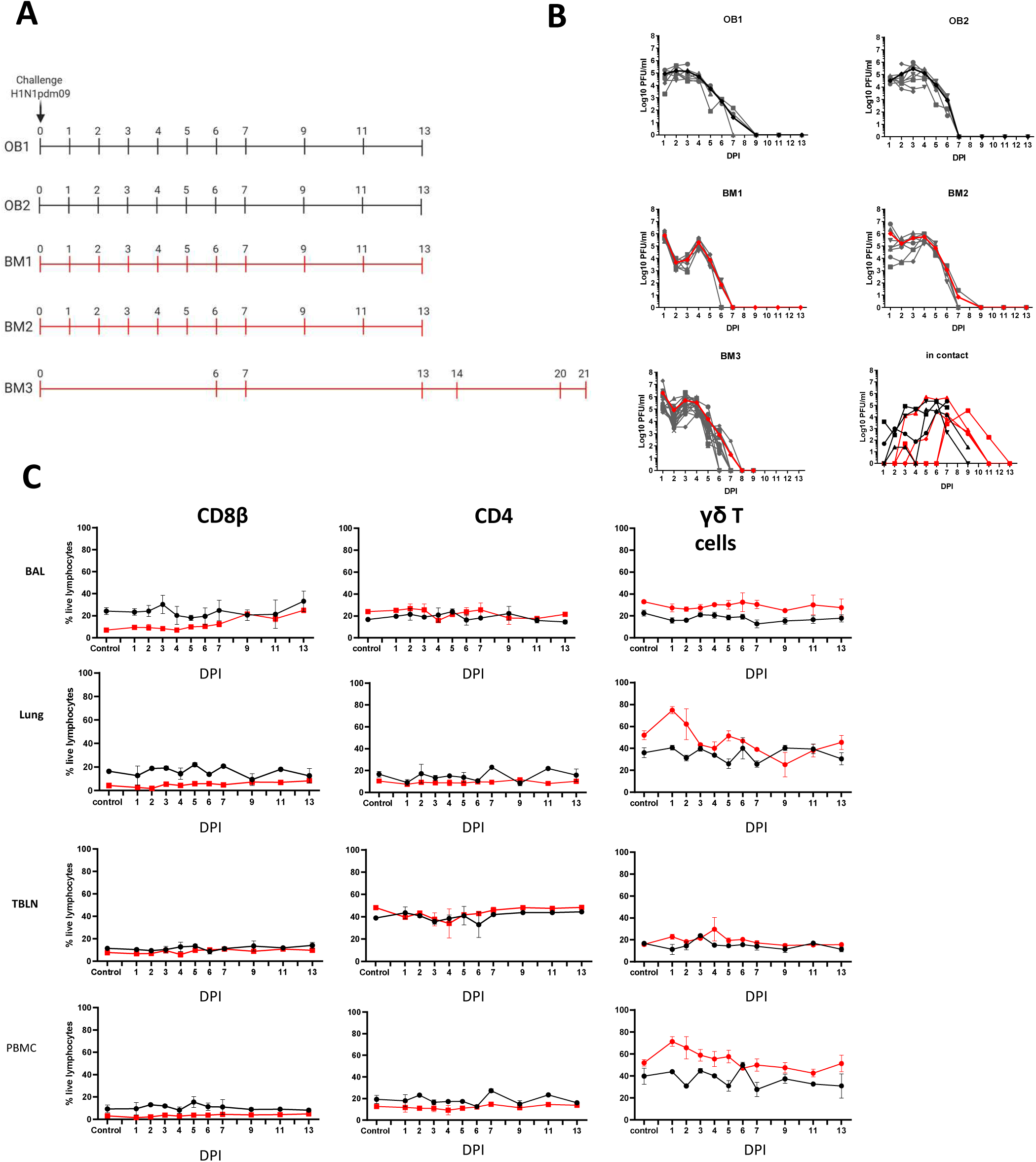
Experimental design, viral load and cell subset dynamics following H1N1 pmd09 infection. (**A)** Pigs were infected with H1N1pdm09 and culled on the days indicated. Two experiments with outbred (OB1 and OB2, black line) and two with inbred Babraham pigs (BM1 and BM2, red line) were performed. Two in-contact animals were included in each experiment one culled at day 11 and one at day 13 post infection. An extended time course of 21 days was performed with 18 inbred Babraham pigs (BM3, red line) with animals culled on the indicated days. (**B)** Virus load was determined by plaque assay of daily nasal swabs at the indicated time points. The thick line indicate the mean. (**C)** Proportions of CD4, CD8 and γδ cells were determined by flow cytometry at the indicated time points.

### Tissue sample and processing

Two nasal swabs (one per nostril) were taken from all surviving pigs following infection with H1N1pdm09 (**Fig. 1**) on days 1 to 7, 9, 11 and 13 in OB1, OB2, BM1 and BM2, and on days 1 to 9 in BM3. Animals were humanely euthanized at the indicated times with an overdose of pentobarbital sodium anaesthetic. Peripheral blood (PBMC), tracheobronchial lymph nodes (TBLN), lung, bronchial alveolar lavage (BAL) were processed as previously described (Morgan et al., 2016b; Holzer et al., 2018b). The tissue homogenate was washed, red blood cells lysed and cell suspension passed through 100μM cell strainer twice. Cells were cryopreserved in FBS containing 10% DMSO.

### Plaque assays

Virus titer in nasal swabs was determined by plaque assay on MDCK cells (Central Service Unit, The Pirbright Institute, UK). Samples were 10-fold serially diluted in Dulbecco’s Modified Eagle’s Medium (DMEM) and 200µl overlayered on confluent MDCK cells in 12 well tissue culture plates. After 1 hour, the plates were washed and overlayered with 2ml of culture medium containing 0.66 % Agar. Plates were incubated at 37°C for 48 to 72 hours and plaques visualized using 0.1% crystal violet. plaques were counted at the appropriate dilution and expressed as plaque forming units (PFU) per ml of nasal swab.

### IFNγ ELISpot assay

Frequencies of IFNγ spot forming cells (SFC) were determined using cryopreserved cells a previously described (Morgan et al., 2016a; Holzer et al., 2018a). Cells were stimulated with live MDCK-grown H1N1pdm09 (MOI 1), medium control, or 4 μg/ml Con A (Sigma-Aldrich). Results were expressed as number of IFNγ producing cells per 10^6^ cells after subtraction of the average number of spots in medium control wells.

### Flow cytometry

Cryopreserved single cell suspensions from blood, TBLN, BAL and lung were thawed, rested for 1-2 hours and aliquoted into 96 well plates at 1×10^6^ cells/ well. Cells were stimulated with live MDCK-grown H1N1pdm09 (MOI 1) or medium control and incubated at 37°C for 18 hours. Golgi plug (BD Biosciences) was added for the last 4 hours of stimulation. PMA Ionomycin (Biolegend) was added to appropriate control wells as a positive control at the same time as the Golgi plug. Following incubation cells were washed at 1000 x g for 5 minutes and re-suspended followed by addition of primary antibodies, Near-Infrared Fixable LIVE/DEAD stain (Invitrogen) and secondary antibodies (**Table 3**). Cells were fixed and permeabilised with BD Fix and perm buffer (BD Biosciences) as per the manufacturer’s instructions prior to the addition of internal cytokine antibodies. Cells were washed and re-suspended in PBS prior to analysis using a MACSquant analyser10 (Miltenyi).

The NP_290-298_ SLA tetramer binding was performed as previously described (Tungatt et al., 2018). Briefly, biotinylated NP peptide loaded SLA monomers, were freshly assembled into tetramer with streptavidin BV421 (Biolegend, UK) and diluted with PBS to a final concentration of 0.1 μg/μl. Two million mononuclear cells were incubated with protease kinase inhibitor (Dasatinib, Axon Medchem) in PBS for 30 minutes at 37°C and 0.3 µg of tetramer was added to the cells on ice for another 30 minutes. Surface staining with optimal antibody concentrations in FACS buffer (PBS supplemented with 2% FCS and 0.05% sodium azide) was performed on ice for 20 minutes (**Table 1**). Samples were washed twice with FACS buffer and fixed in 1% paraformaldehyde before analysis on MACSquant analyser10 (Miltenyi). All flow cytometry data was analysed by Boolean gating using FlowJo v10.6 (TreeStar, US).

**Table 1.** Comparison of different populations of T cells from Babraham pigs infected with H1N1pdm09 virus.

| % T cells binding to tetramer or /cytokine producing |  | Days Post Infection (DPI) |  |  |
| --- | --- | --- | --- | --- |
|  |  | 6 DPI | 7DPI | 13 DPI |
| **CD8β** | NP<sub>290-298</sub> | no significant (P>0.05) differences* | BAL†>PBMC (P=0.01)<br>lung>PBMC (P=0.04)<br>lung>TBLN (P=0.04) | BAL>lung (P=0.04)<br>BAL>TBLN (P=0.01)<br>BAL>PMBC (P=0.01)<br>lung>PMBC (P=0.01)<br>TBLN>PMBC (P=0.01) |
| **CD8β** | IFNγ | BAL>PBMC (P=0.01)<br>BAL>TBLN (P=0.01)<br>lung>PBMC (P=0.01)<br>lung>TBLN (P=0.03) | BAL>PBMC (P=0.01)<br>BAL>TBLN (P=0.01)<br>lung>PBMC (P=0.01)<br>lung>TBLN (P=0.03) | BAL>PBMC (P=0.01)<br>BAL>TBLN (P=0.01)<br>lung>PBMC (P=0.01)<br>lung>TBLN (P=0.03) |
|  | IL-2 | no significant (P>0.05) differences | BAL> lung (P=0.06)<br>BAL>PBMC (P=0.02)<br>BAL>BM TBLN (P=0.02) | BAL>lung (P=0.01)<br>BAL>PBMC (P=0.01)<br>BAL>TBLN (P=0.01) |
|  | TNF | BAL>lung (P=0.01)<br>BAL>PBMC (P=0.01)<br>BAL>TBLN (P=0.01)<br>lung>PBMC (P=0.02) | BAL>lung (P=0.02)<br>BAL>PBMC (P=0.01)<br>BAL>TBLN (P=0.01)<br>lung>PBMC (P=0.02) | BAL>lung (P=0.01)<br>BAL>PBMC (P=0.01)<br>BAL>TBLN (P=0.01)<br>lung>PBMC (P=0.02)<br>lung>TBLN (P=0.06) |
| **CD4** | IFNγ | no significant (P>0.05) differences | lung>TBLN (P=0.02)<br>lung>PBMC (P=0.01) | BAL>PBMC (P=0.06)<br>lung>PBMC (P=0.06) |
|  | IL-2 | BAL>PBMC (P=0.01)<br>lung>PBMC (P=0.03)<br>TBLN>PBMC (P=0.01) | BAL>PBMC (P=0.01)<br>lung>PBMC (P=0.01)<br>TBLN>PBMC (P=0.01) | BAL>PBMC (P=0.01)<br>lung>PBMC (P=0.01)<br>TBLN>PBMC (P=0.01) |
|  | TNF | no significant (P>0.05) differences | no significant (P>0.05) differences | BAL>PBMC (P=0.06)<br>TBLN> lung (P=0.06)<br>TBLN>PBMC (P=0.02) |

\* P-values based on pairwise Mann-Whitney-Wilcoxon tests following a significant (P<0.05) Kruskal-Wallis test
† BAL: broncho-alveolar lavage; PBMC: peripheral blood mononuclear cell; TBLN: tracheobronchial lymph node

### Serological assays

ELISA was performed using live H1N1pdm09 virus or recombinant haemagglutinin from H1N1pdm09 (pH1) containing a C-terminal thrombin cleavage site, a trimerization sequence, a hexahistidine tag and a BirA recognition sequence as previously described (Huang et al., 2015). Cut-off values determined as average naïve values plus three-fold standard deviation at optimal starting dilution. Starting dilutions were 1:20, 1:2 and 1:4 for serum, BAL and nasal swab respectively. Hemagglutination inhibition (HAI) Ab titers were determined using 0.5% chicken red blood cells and H1N1pdm09 at a concentration of 4 HA units/ml. Microneutralization (MN) was performed using standard procedures as described previously (Powell et al., 2012; McNee et al., 2020).

The porcine sera were also tested for binding to MDCK-SIAT1 cells stably expressing pH1 from H1N1pdm09 (A/England/195/2009), H1 from A/Puerto Rico/8/1934 (PR8, H1N1) and H5 HA (A/Vietnam/1203/2004). Confluent cell monolayers in 96-well microtiter plates were washed with PBS and 50 μl of the serum dilution was added for 1 h at room temperature. The plates were washed three times with PBS and 100 μl of horseradish peroxidase (HRP)-conjugated goat anti-pig Fc fragment secondary antibody (Bethyl Laboratories, diluted in PBS, 0.1% BSA) was added for 1 h at room temperature. The plates were washed three times with PBS and developed with 100 µl/well TMB high sensitivity substrate solution (Biolegend). After 5 to 10 min the reaction was stopped with 100 µl 1M sulfuric acid and the plates were read at 450 and 570 nm with the Cytation3 Imaging Reader (Biotek). The cut off value was defined as the average of all blank wells plus three times the standard deviation of the blank wells.

### Enzyme-linked lectin Assay (ELLA)

Neuraminidase inhibiting Ab titres were determined in serum and BAL fluid using an Enzyme-linked lectin assay (ELLA). Ninety six-well microtiter plates (Maxi Sorp, Nunc, Sigma-Aldrich, UK) were coated with 50 μl/well of 25 μg/ml fetuin and incubated at 4°C overnight. Heat inactivated sera samples were serially diluted in a separate 96-well plate. An equal volume of (H7(Net219) N1(Eng195) S-FLU (H7N1 S-FLU) (kindly provided by Professor Alain Townsend, University of Oxford) was added to each well and incubated at room temperature on a rocking platform for 20 minutes. The H7N1 S-FLU was titered beforehand in the absence of serum to determine optimal concentration for the assay. Fetuin plates were washed with PBS four times before 100 μl/well of the serum/virus mix was transferred and incubated overnight at 37°C. The serum/virus mix was removed, and the plate washed four times with PBS before adding 50μl/well of Peanut Agglutinin-HRP at 1 μg/ml and incubating for 90 minutes at room temperature on a rocking platform. Plates were washed and 50 μl/well of TMB High Sensitivity substrate solution (BioLegend, UK) was added. Plates were developed for 6 minutes, the reaction stopped with 50 μl of 1M H_2_SO_4_ and the plates were read at 450 and 630 nm using a Biotek Elx808 reader. Samples were measured as end titre representing the highest dilution with signal greater than cut-off. The cut off value was defined as the average of all blank wells plus three times the standard deviation of the blank wells.

### B cell ELISpot

B cell ELISpots were performed for the detection and enumeration of antibody-secreting cells in single cell suspensions prepared from different tissues and peripheral blood. ELISpot plates (Multi Screen-HA, Millipore, UK) were coated with 100 µl per well of appropriate antigen or antibody diluted in carbonate/bicarbonate buffer for 2h at 37°C. To detect HA-specific spot-forming cells, plates were coated with 2.5 µg per well of recombinant pHA from H1N1pdm09 (A/England/195/2009) and for the enumeration of total IgG-secreting cells with 1 µg per well of anti-porcine IgG (mAb, MT421, Mabtech AB, Sweden) or with culture medium supplemented with 10% FBS (media background control). The coated plates were washed with PBS and blocked with 200 µl/well 4% milk (Marvel) in PBS. Frozen cell suspensions from different tissues were filtered through sterile 70 µM cell strainers, plated at different cell densities in culture medium (RPMI, 10% FBS, HEPES, Sodium pyruvate, Glutamax and Penicillin/Streptomycin) on the ELISPOT plates and incubated for a minimum of 18 h at 37°C in a 5% CO_2_ incubator. After incubation the cell suspension was removed, the plates washed once with ice-cold sterile H_2_O and thereafter with PBS/0.05 % Tween 20, before incubation with 100 µl per well of 0.5 µg/ml biotinylated anti porcine IgG (mAb, MT424, Mabtech AB, Sweden) diluted in PBS/0.5 % FBS for two hours at room temperature. Plates were washed with PBS/0.05% Tween 20 and incubated with streptavidin – alkaline phosphatase conjugate (Strep-ALP, Mabtech AB, Sweden). After a final wash, the plates were incubated with AP Conjugate Substrate (Bio-Rad, UK) for a maximum of 30 min. The reaction was stopped by rinsing the plates in tap water and dried before spots were counted.

### Statistical analysis

All statistical analysis was performed using Prism 8.1.2. The kinetics of viral shedding were analysed using a linear mixed model. The model included viral titre (log_10_ PFU/ml) as the response variable, day post infection (as a categorical variable) and pig type (OB or BM) and an interaction between them as fixed effects and pig ID nested in experiment as random effects. The model was implemented using the lme4 package (Bates et al., 2015) in R (version 3.6.1) ((https://www.R-project.org/)

ELISpot data were analysed using a linear model. The model included log_10_ SFC/10^6^ cells+1 as the response variable and day post infection (as a categorical variable), source (BAL, lung, PBMC, TBLN) and pig type (OB or BM) and two- and three-way interactions between them as fixed effects. Model simplification proceeded by stepwise deletion of non-significant (P>0.05) terms as judged by *F*-tests. The model was implemented in R (version 3.6.1).

Because of possible non-normality and non-constant variance the percentage of different T cells (NP_290-298_ CD8, IFNγ CD8β, IL-2 CD8β, TNF CD8β, IFNγ CD4, IL-2 CD4, TNF CD4, IFNγ CD2 γδ ex vivo, TNF CD2 γδ ex vivo, IFNγ/TNF CD2 γδ ex vivo, IFNγ CD2 γδ, TNF CD2 γδ, IL-17A CD2 γδ) from each source (BAL, lung, PBMC, TBLN) and pig type (OB and BM) at each day post infection were analysed using Kruskal-Wallis tests. If significant (P<0.05), pairwise Mann-Whitney-Wilcoxon tests were used to compare groups. These analyses were implemented in R (version 3.6.1). A similar approach was used to compare the percentage of different T cells in all sources from inoculated and uninfected control pigs at each time point and in BAL from in-contact and experimentally-inoculated pigs (in this case observations from 6-11 dpi were combined).

The dynamics of antibody responses were analysed by fitting logistic growth curves to the data, *y*=*κ*/(1+exp(-*β*(*t*-*δ*)), where *y* is the log_10_ antibody titre, *t* is days post infection, *κ* is the upper asymptote, *β* is the rate of increase and *δ* is the time of maximum increase. The parameters (i.e. *κ, β* and *δ*) were allowed to vary between BM and OB pigs. Model fitting using the nlme package (https://CRAN.R-project.org/package=nlme) in R (version 3.6.1).

## Results

### Experimental design, virus shedding and lymphocyte dynamics during H1N1pdm09 infection

Five experiments were performed to characterise local and systemic immune responses. In the first four experiments ten pigs were infected intranasally with H1N1pdm09 virus and monitored for clinical signs. One infected pig was culled on each of days 1 to 7, 9, 11 and 13 post infection. A full post-mortem examination was performed and BAL, lung, TBLN and PBMC samples collected. Four uninfected controls were sampled in parallel, two on the day prior to infection and two at day 8 post infection. Two experiments with outbred (OB) pigs (referred to as experiments OB1 and OB2) and two with inbred Babraham (BM) pigs (referred to as BM1 and BM2) were performed (**Fig. 1A**). In addition, 2 naïve pigs (referred to as in-contact pigs) were co-housed with the directly challenged pigs in experiments OB1, OB2, BM1, BM2 and culled at days 11 and 13 post contact. A fifth experiment was carried out with 18 BM (experiment BM3) in which 3 pigs were culled on each of days 6, 7, 13,14, 20 and 21 post infection (**Fig. 1A**). In the BM3 experiment six uninfected controls were sampled, 3 one day before and 3 on the day of infection.

Viral load was determined in daily nasal swabs taken from both the directly challenged and in-contact pigs (**Fig. 1B)**. In directly challenged pigs, peak virus load was reached 1 to 3 days post infection (DPI), declined sharply after 4 DPI and was not detectable after 7 DPI. No differences in virus shedding between OB and BM were detected (p=0.65). Although the onset of viral shedding was delayed, most in-contact pigs showed similar kinetics to directly challenged ones, indicating that the natural contact infection is very similar to intra-nasal challenge with mucosal atomization device (MAD).

We determined the proportion of CD8β, CD4 and γδ cells over the time course in BAL, lung, TBLN and PBMC (**Fig. 1C, Suppl Fig1**). BM animals had a significantly lower proportion of CD8β T cells than OB, apparent in all tissues in naïve unexposed animals (6.6% in BM vs 24.2% in OB in BAL, 4.2% vs 16.2% in lung, 3.2% vs 9.4% in PBMC and 7.6% vs 11.5% in TBLN) (**Suppl Fig. 1A**). BM animals also showed a significantly higher proportion of γδ T cells in BAL, lung and PBMC (**Suppl Figure 1A**). No significant differences in CD4 T cells were detected between OB and BM. The proportion of CD4, CD8 and γδ cells did not change significantly over the time course of H1N1pdm09 infection, although an increase in the proportion of CD8β in the BAL for the BM animals was observed, as previously reported (Khatri et al., 2010).

Overall the kinetic of virus infection and shedding were similar between BM and OB, although there were differences in the proportions of proportions of CD8 and γδ cells.

### T cell responses during H1N1pdm09 infection in pigs

As T cells are crucial for control of virus replication, we examined in detail the CD8 and CD4 responses during H1N1pdm09 infection (McMichael et al., 1983; Sridhar et al., 2013; Hayward et al., 2015; Holzer et al., 2019). First, we enumerated IFNγ secreting cell by ELISpot following re-stimulation with H1N1pdm09 (**Figs. 2A and B**). IFNγ spot forming cells (SFC) were detectable from 6 DPI and maintained in all tissues until 21 DPI. During the early stage of infection the strongest responses were in the TBLN (mean 474 SFC/10^6^ cells at 7 DPI), whereas from 14 to 21 DPI the highest number of IFNγ secreting cells was detected in the lung, with SFC continuing to expand in this tissue (mean 368 SFC/10^6^ cells at 14 DPI and 972/10^6^ cells SFC at 21 DPI). The response in the BAL was lower than lung (p=0.04), due to the low proportion of T cells present in the BAL (**Fig. 1C**). The IFNγ ELISpot response in the PBMC was low with a peak of 296 SFC/10^6^ cells at 13 DPI. No differences in responses between the same tissues in OB and BM were detected (p>0.11).

**Figure 2.**
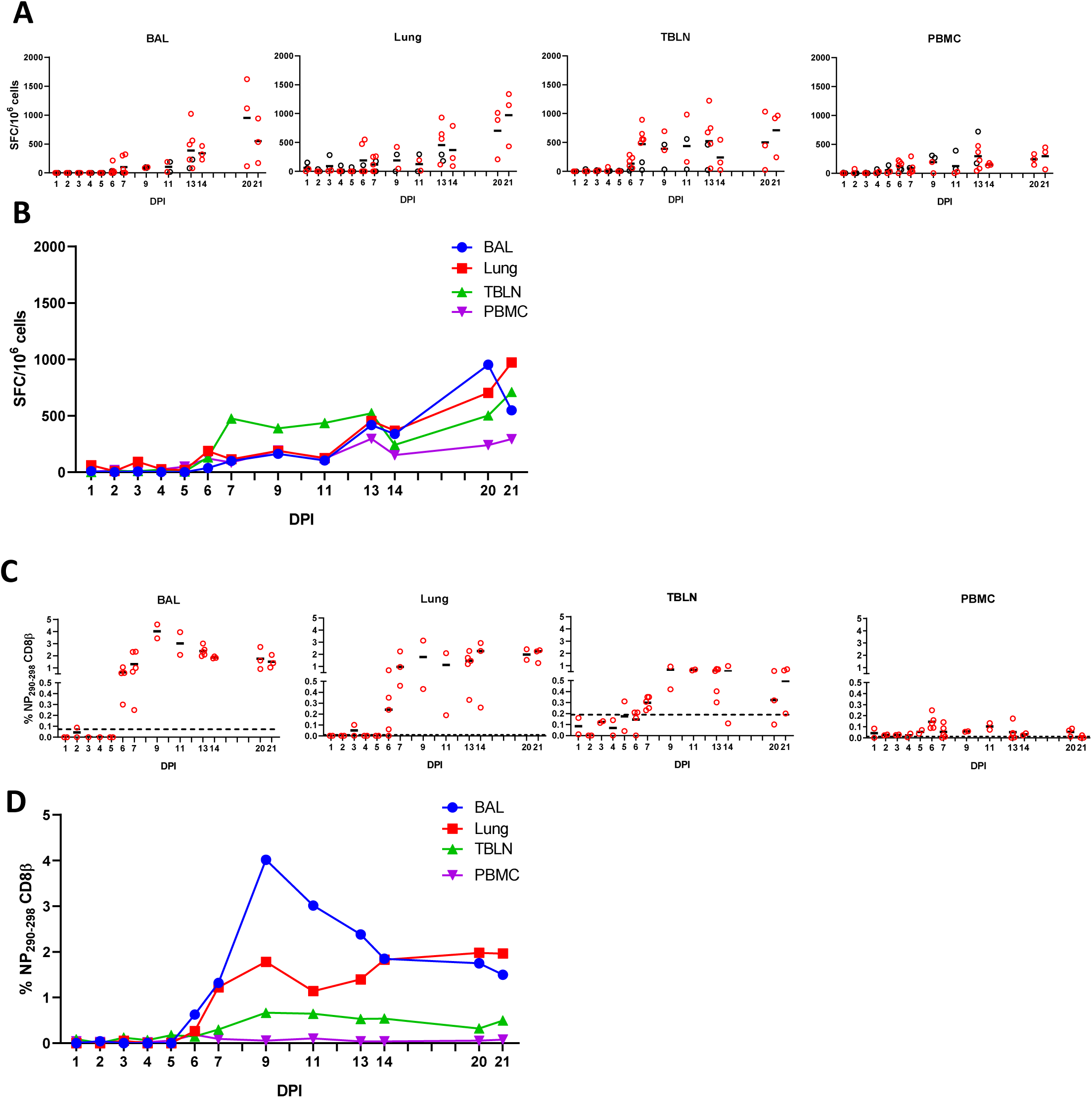
IFNγ ELISpot and tetramer responses. **(A**,**B)** IFNγ secreting spot forming cells (SFC) in BAL, lung, TBLN and PBMC in outbred (black circles) and inbred (red circles) pigs were enumerated after stimulation with H1N1pdm09 or medium control. **(B)** The mean percentages for each population are shown. DPI 1 to 7, 9, 11 and 13 each show results from 2 outbred and 2 inbred pigs. DPI 6, 7, 13, 14, 20 and 21 also include results from 3 additional inbred pigs. **(C, D)** Proportions of NP_290-298_ CD8 T cells in tissues. Background staining with SLA matched tetramers containing irrelevant peptide has been subtracted. Dotted lines indicates proportions of tetramer positive cells in uninfected animals. Data from 2 outbred and 2 inbred pigs is shown for days 1 to 5 and 9 to 11. DPI 6, 7, 13, 14, 20 and 21 show data from 3 additional inbred pigs. Data in C and D is from 2 pigs (DPI 1-5, 9, 11), 3 pigs (DPI 14, 20, 21) or 5 pigs (DPI 6, 7 and 13).

To further dissect the T cell response, we enumerated antigen specific cytotoxic CD8β T cells against the nuclear protein (NP) using peptide NP_290-298_ (DFEREGYSL) tetramer, which we have previously shown to be dominant in BM animals infected with H1N1pdm09 (Tungatt et al., 2018). Tetramer responses were measured in experiments BM1, BM2 and BM3 (**Suppl Fig 1B, Figs. 2C and D**). NP_290-298_ responses were detected in BAL and lung at 6 DPI, reaching a peak at 9 DPI and still present at 20 - 21 DPI. In TBLN one animal responded at 5 DPI, but the peak was at 9 - 11DPI and still present at 21 DPI. The responses in PBMC were low (0.2% at 6 DPI) and there were no detectable responses at 20-21 DPI **(Table 1)**.

### Cytokine production by CD4 and CD8 T cells

We analysed production of IFNγ, TNF and IL-2 by CD8β and CD4 cells by intracellular staining (ICS) (**Suppl Fig. 2A**). The kinetics of the CD8 cytotoxic T cell response was similar when analysed by ICS, ELISpot and tetramer binding. There was a minimal response up to 5 - 6 DPI, followed by a marked increase in cytokine-producing T cells particularly in the BAL (peak of 7.9% IFNγ and 7.6% TNF at 9 DPI) and lung (peak of 1.3% IFNγ and 0.6% TNF at 9 DPI). CD8 T-cells produced minimal IL-2 in all tissues except for BAL, where 0.7% to 1.3% positive cells were detected between 7-13 DPI. PBMC had much lower proportion of cytokine producing CD8 cells with maximum 0.3% IFNγ and 0.2% TNF production in PBMC at 9 DPI. The high cytokine responses were maintained in BAL and lung until 21 DPI, with lower responses in the TBLN and none in PBMC (**Fig. 3**).

**Figure 3.**
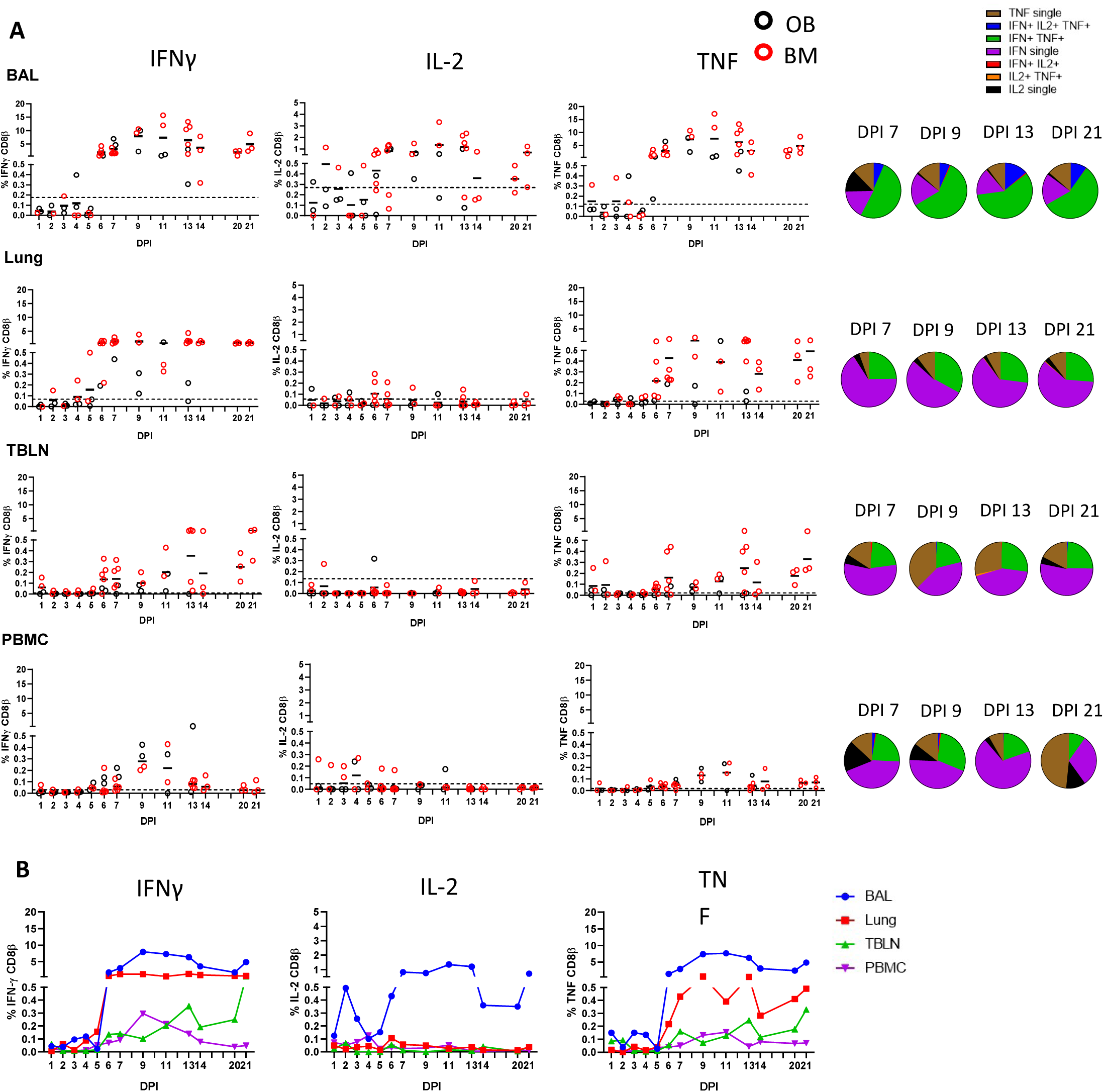
CD8β T cell cytokine responses. **(A)** Cytokine response of CD8β T cells in outbred (black circles) and inbred (red circles) pigs at each time point following influenza infection. BAL, lung, TBLN or PBMC cells were stimulated with H1N1pdm09 and cytokine secretion measured using intra-cytoplasmic staining. The mean of the 22 uninfected control animals is represented by a dotted line. DPI 1 to 7, 9, 11 and 13 each show results from 2 outbred and 2 inbred pigs. DPI 6, 7, 13, 14, 20 and 21 also include results from 3 additional inbred pigs. Pie charts show the proportion of single, double and triple cytokine secreting CD8 T cells for IFNγ, TNF and IL-2 at 7, 9, 13 and 21 DPI. **(B)** The mean percentages for IFNγ, TNF and IL-2 in each tissue for both OB and BM together are shown over the time course.

We next determined the quality of cytokine responses of CD8β cells. The CD8β T-cell cytokine response was dominated by IFNγ single producing cells with some IFNγ/TNF double producing cells also present in all tissues (**Fig. 3A**). However, the highest proportion of double IFNγ/TNF producing cells was present in the BAL **(Table 1**). A triple secreting IFNγ/ IL-2/TNF population was detected only in the BAL and these cells produced greater levels of IFNγ per cell as measured by MFI (**Suppl Fig. 2B**). The individual cytokine profiles of the BM and OB were similar during the time course of H1N1pdm09 infection and shown in **Suppl Fig. 3**. We also analysed the responses in the in-contact animals from experiments OB1, OB2, BM1 and BM2. These animals had the same profiles of cytokine production in BAL (**Suppl Fig. 4**) and in the other tissues (data not shown) as directly challenged animals (**Table 2**).

**Table 2.** Comparison of T cells in broncho-alveolar lavage from experimentally-inoculated (I) and in-contact (C) Babraham (BM) and outbred (OB) pigs infected with H1N1pdm09 swine influenza virus.

| T cells | % cells producing |  |  |
| --- | --- | --- | --- |
| | IFNγ | TNF | IL-2 or IL-17a* |
| CD8β | no significant<br>(P>0.05) differences† | no significant<br>(P>0.05) differences† | no significant<br>(P>0.05) differences† |
| CD4 | no significant<br>(P>0.05) differences† | OB, C>BM, C<br>(P=0.03)<br>OB, C>BM, I<br>(P=0.02) | no significant<br>(P>0.05) differences† |
| $\gamma\delta$ *ex vivo* | no significant<br>(P>0.05) differences† | no significant<br>(P>0.05) differences† | not tested |
| $\gamma\delta$ H1N1pdm09<br>stimulated | no significant<br>(P>0.05) differences† | no significant<br>(P>0.05) differences† | no significant<br>(P>0.05) differences† |

\* IL-2 for CD8β and CD4 T cells; IL-17a for $\gamma\delta$ T cells
† P-values based on pairwise Mann-Whitney-Wilcoxon tests following a significant (P<0.05) Kruskal-Wallis test

**Table 3.** Antibodies used.

| Antigen | Clone | Isotype | Fluorochrome | Source of primary Ab | Details of secondary Ab |
| --- | --- | --- | --- | --- | --- |
| **Staining for conventional T cells** |  |  |  |  |  |
| CD4 | 74-12-4 | IgG2b | PerCP-Cy5.5 | BD Biosciences |  |
| CD8b | PPT23 | IgG1 | FITC | Bio-Rad Laboratories |  |
| TNF | MAb11 | IgG1 | BV421 | BioLegend |  |
| IFNγ | P2G10 | IgG1 | APC | BD Biosciences | |
| IL-2 | A150D 3F1 2H2 | IgG2a | PE-Cy7 | ThermoFisher | rat-anti-mouse, IgG2a, BioLegend |
| **Staining for $\gamma\delta$ T cells** |  |  |  |  |  |
| TCR $\gamma\delta$ | PGBL22A | IgG1 | PE-Cy7 | Cambridge bioscience | rat-anti-mouse, IgG1, BioLegend |
| CD8α | 76-2-11 | IgG2a | FITC | BD Biosciences | |
| CD2 | MSA4 | IgG2a | PerCP-Cy5.5 | Cambridge bioscience | rat-anti-mouse, IgG2a, BioLegend |
| TNF | MAb11 | IgG1 | BV421 | BioLegend |  |
| IFNγ | P2G10 | IgG1 | APC | BD Biosciences | |
| IL-17A | SCPL1362 | IgG1 | PE | BD Biosciences |  |

**Figure 4.**
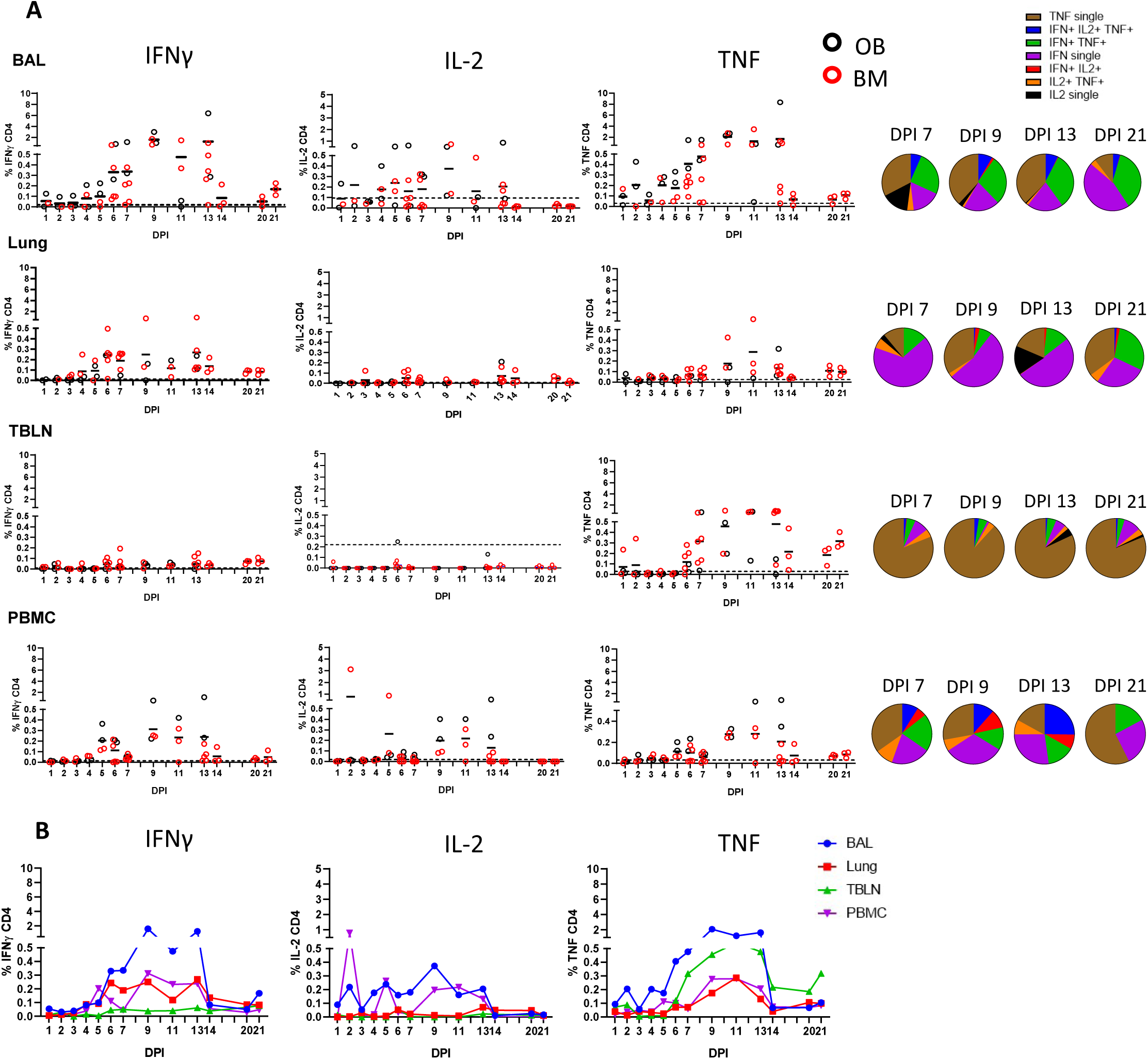
CD4 T cell cytokine responses. **(A)** Cytokine response of CD4 T cells in outbred (black circles) and inbred (red circles) pigs at each time point following influenza infection. BAL, lung, TBLN or PBMC cells were stimulated with H1N1pdm09 and cytokine secretion measured using intra-cytoplasmic staining. The mean of the 22 uninfected control animals is represented by a dotted line. DPI 1 to 7, 9, 11 and 13 each show results from 2 outbred and 2 inbred pigs. DPI 6, 7, 13, 14, 20 and 21 also include results from 3 additional inbred pigs. Pie charts show the proportion of single, double and triple cytokine secreting CD8 T cells for IFNγ, TNF and IL-2 at 7, 9, 13 and 21 DPI. **(B)** The mean percentages for IFNγ, TNF and IL-2 in each tissue for OB and BM together are shown over the time course.

The CD4 response was lower than the CD8 and developed earlier at 4 −5 DPI in some animals (**Fig. 4**). It was greatest in the BAL and peaked at 9 DPI similarly to CD8 (1.6% IFNγ and 2.1% TNF) and almost disappeared by 21 DPI. The CD4 response was lower in TBLN, lung and PBMC (**Table 1**). CD4 cytokine secretion differed between tissues. Single cytokine-secreting IFNγ and TNF CD4 cells were dominant in the lung and TBLN respectively, while in the BAL and PBMC both single IFNγ, single TNF and double IFNγ/TNF were present. The individual cytokine profiles of the BM and OB animals were comparable (**Suppl Fig. 3**). The in-contacts also showed a similar pattern of cytokine production except for TNF (**Suppl Fig. 4, Table 2**).

These results demonstrate that there is a strong antigen specific CD8 response in the local lung tissues and in particularly in the BAL. Cytokine production by CD8 was dominated by IFNγ and TNF, but the BAL also had a significant population of IL-2-producing cells and more double- and triple-producing cells, compared to TBLN, lung and PBMC. The CD4 T-cell response was also greatest in the BAL, although much lower and declining more rapidly than the CD8 response. The cytokine responses were similar between the in-contact and directly infected animals, indicating the similarities between experimental intra-nasal challenge and natural infection. No differences in magnitude, kinetic and quality of cytokine responses were observed between the OB and BM animals.

### γδ T cell responses during H1N1pdm09 infection in pigs

The importance of γδ T cells in control of influenza infection has been demonstrated in mice and humans (Carding et al., 1990; Li et al., 2013a; Xue et al., 2017; Palomino-Segura et al., 2020). In pigs γδ T cells are a prominent population in blood and secondary lymphoid organs and can produce IFNγ, TNF and IL-17A following polyclonal stimulation (Takamatsu et al., 2006; Gerner et al., 2009; Sedlak et al., 2014). Porcine γδ T cells have been divided into different subsets based on the expression of CD2 and CD8α (Stepanova and Sinkora, 2013).

We measured IFNγ, TNF and IL-17A production in CD2^+^ γδ cells immediately *ex vivo* and following H1N1pdm09 re-stimulation. The cytokine production by CD2^-^ γδ T cells was very low (data not shown) and thus we focused on CD2^+^ γδ T cells. *Ex vivo*, BAL CD2^+^ γδ cells, without H1N1pdm09 stimulation, secreted IFNγ and TNF early post infection with the highest frequency of 0.6% IFNγ and 1.6% TNF at 3 DPI. The cells co-produced IFNγ/TNF at low levels **(Fig. 5)**. A minimal amount of IL-17A was detected in BAL and no IFNγ, TNF or IL-17 in the other tissues *ex vivo* (data not shown).

**Figure 5.**
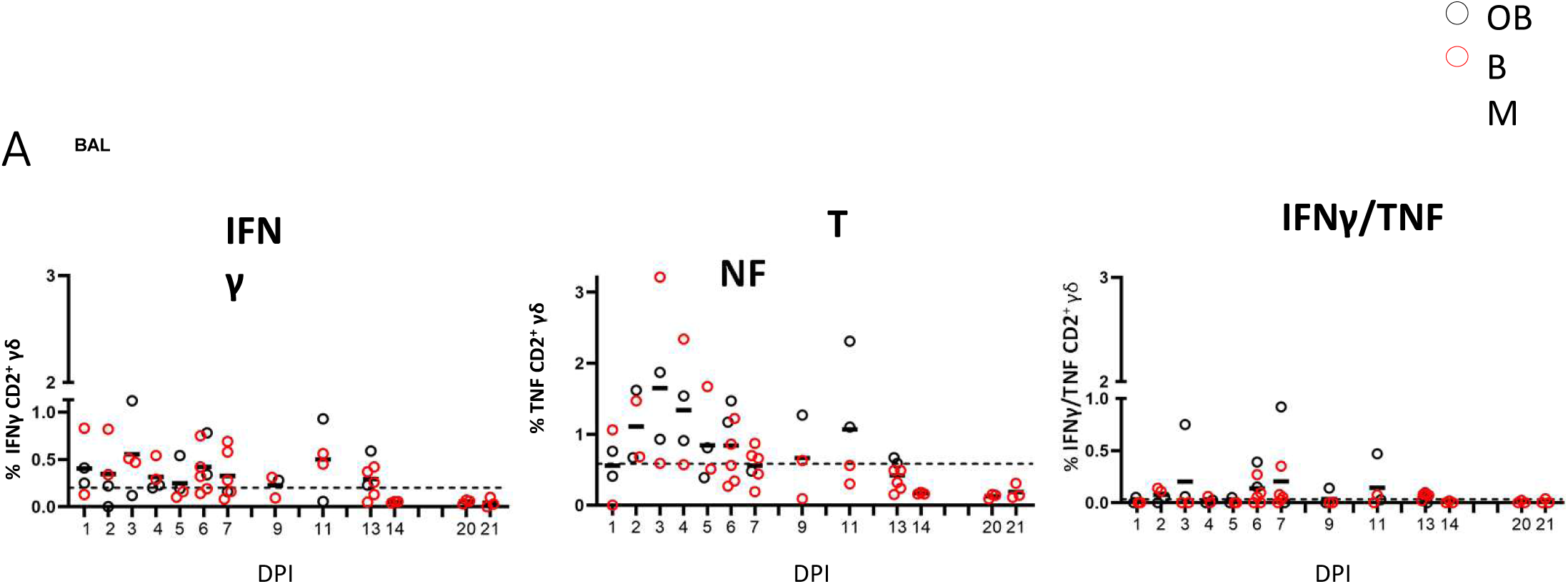
γ δ T cell *ex vivo* responses in BAL. Cytokine response of γ δ T cells in outbred (black circles) and inbred (red circles) pigs at each time point following H1N1pdm09 infection. IFNγ and TNF production in BAL cells *ex vivo* without stimulation was measured using intra-cytoplasmic staining. The right hand panel shows the IFNγ/TNF double producing cells. The mean of the 22 uninfected control animals is represented by a dotted line. DPI 1 to 7, 9, 11 and 13 each show results from 2 outbred and 2 inbred pigs. DPI 6, 7, 13, 14, 20 and 21 also include results from 3 additional inbred pigs.

We also measured cytokine production after H1N1pdm09 stimulation *in vitro* and the highest proportion of IFNγ producing cells was detected in BAL and lung at 7 DPI and maintained until 13 DPI (0.4% at 11 DPI for BAL) while in lung the highest frequency was 0.4% at 7 DPI (**Fig. 6**). TNF showed similar pattern in BAL: increased at 9 DPI reaching a peak at 11 DPI with a mean of 0.7%. At later stages of infection, the frequency of cytokine producing cells were much lower. The majority of the H1N1pdm09 stimulated cells were IFNγ/TNF co-producing (**Suppl Fig. 5)**. The responses in the contacts (0.5% for IFNγ and more than1 % for TNF) were similar to those in the directly challenged animals following H1N1pmd09 stimulation (**Table 2**). The proportion of IL-17A secreting CD2^+^ cells was much lower compared to IFNγ and TNF with the greatest response in the BAL at 11 DPI (0.2%).

**Figure 6.**
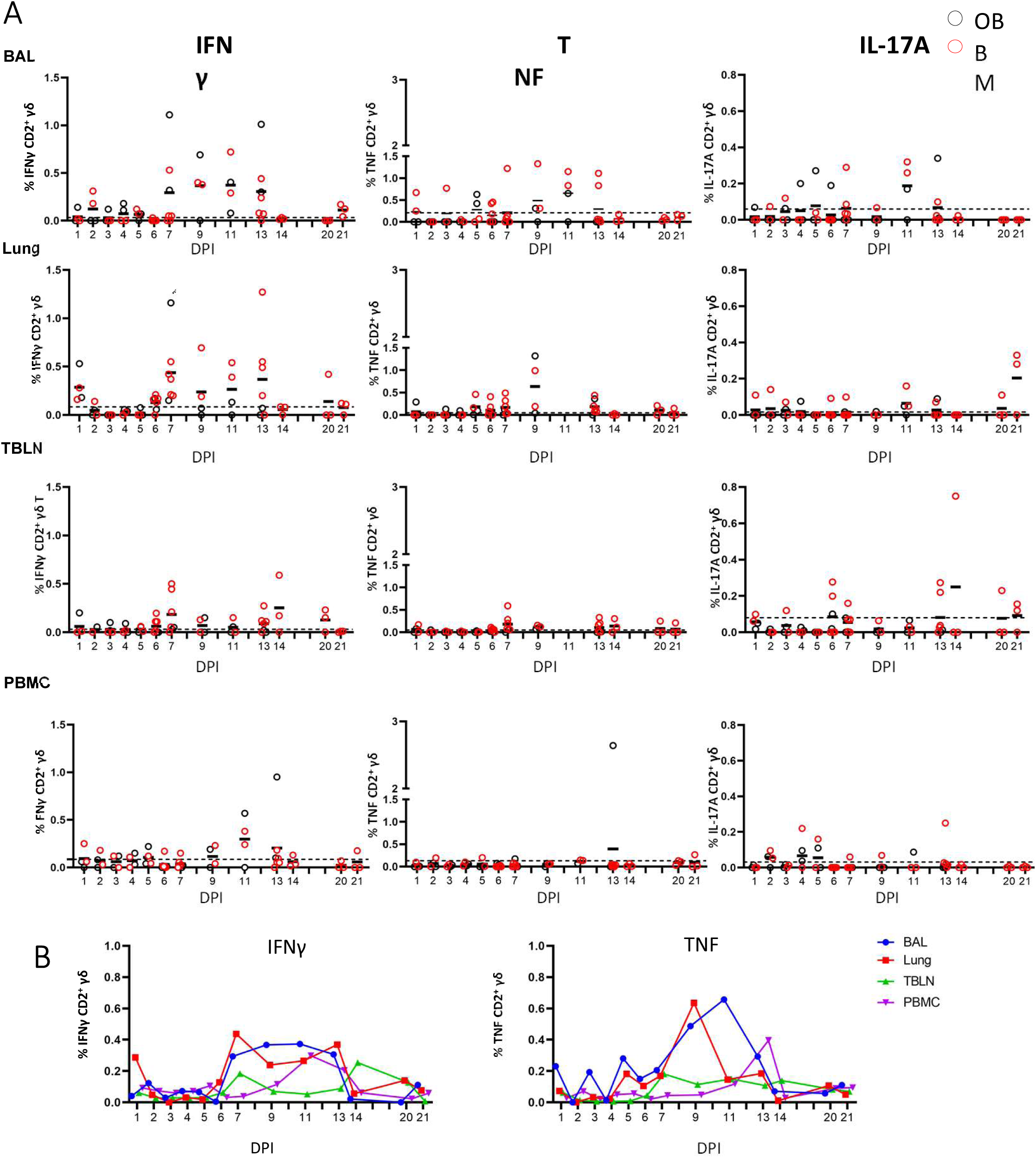
γδ T cell responses after H1N1pdm09 stimulation. **(A)** Frequencies of IFNγ, TNF and IL-17A producing CD2^+^ γδ T cells in outbred (black circles) and inbred (red circles) pigs following influenza infection. BAL, lung, TBLN and PBMC were stimulated with H1N1pdm09 and cytokine secretion measured using intra-cytoplasmic staining. The mean of the 22 uninfected control animals is represented by a dotted line. DPI 1 to 7, 9, 11 and 13 each show results from 2 outbred and 2 inbred pigs. DPI 6, 7, 13, 14, 20 and 21 also include results from 3 additional inbred pigs. **(B)** Mean percentages for IFNγ and TNF in each tissue are shown over the time course.

Overall these data demonstrate that γδ cells produce cytokines *ex vivo* early post infection, but that H1N1pdm09 *in vitro* stimulation increases cytokine production in CD2^+^ γδ T cells from 7 to 13 DPI. No difference between OB and BM pigs were detected of response of *ex vivo* or stimulated γδ cells.

### Antibody and B cell responses during H1N1pdm09 infection in pigs

The antibody response after H1N1pdm09 infection was determined in serum, BAL and nasal swabs. Virus specific IgG and IgA were measured by end point titer ELISA against H1N1pdm09 virus or recombinant HA from H1N1pdm09/A/England/195/2009 (pH1) (**Figs. 7A and B**). Serum IgG against H1N1pdm09 virus was detectable at 5-6 DPI, reached its peak at 14 DPI (1:13,650) and was maintained until 21 DPI (1:8,530). IgA titers were lower compared to IgG. In contrast in BAL, IgG and IgA against H1N1pdm09 were present at the same levels. BAL IgG reached a peak of 1:2,370 at 13 DPI which was maintained up to 21 DPI. IgG and IgA were also measured in nasal swabs from experiment BM3 up to 9 DPI. Responses were detected at 6 DPI reaching a peak of 1:48 and 1:28 respectively by 9 DPI. We measured the ELISA response to pH1, which had a similar kinetic as the response to H1N1pdm09 virus but with approximately a log lower titer (**Fig. 7B)**. No significant differences in the upper asymptote, rate of increase in titre or time of maximum increase were detected for IgG or IgA between OB and BM, except for serum IgA H1N1 ELISA (upper asymptote OB 1:2,700 > BM 1:2,100, p=0.05).

**Figure 7.**
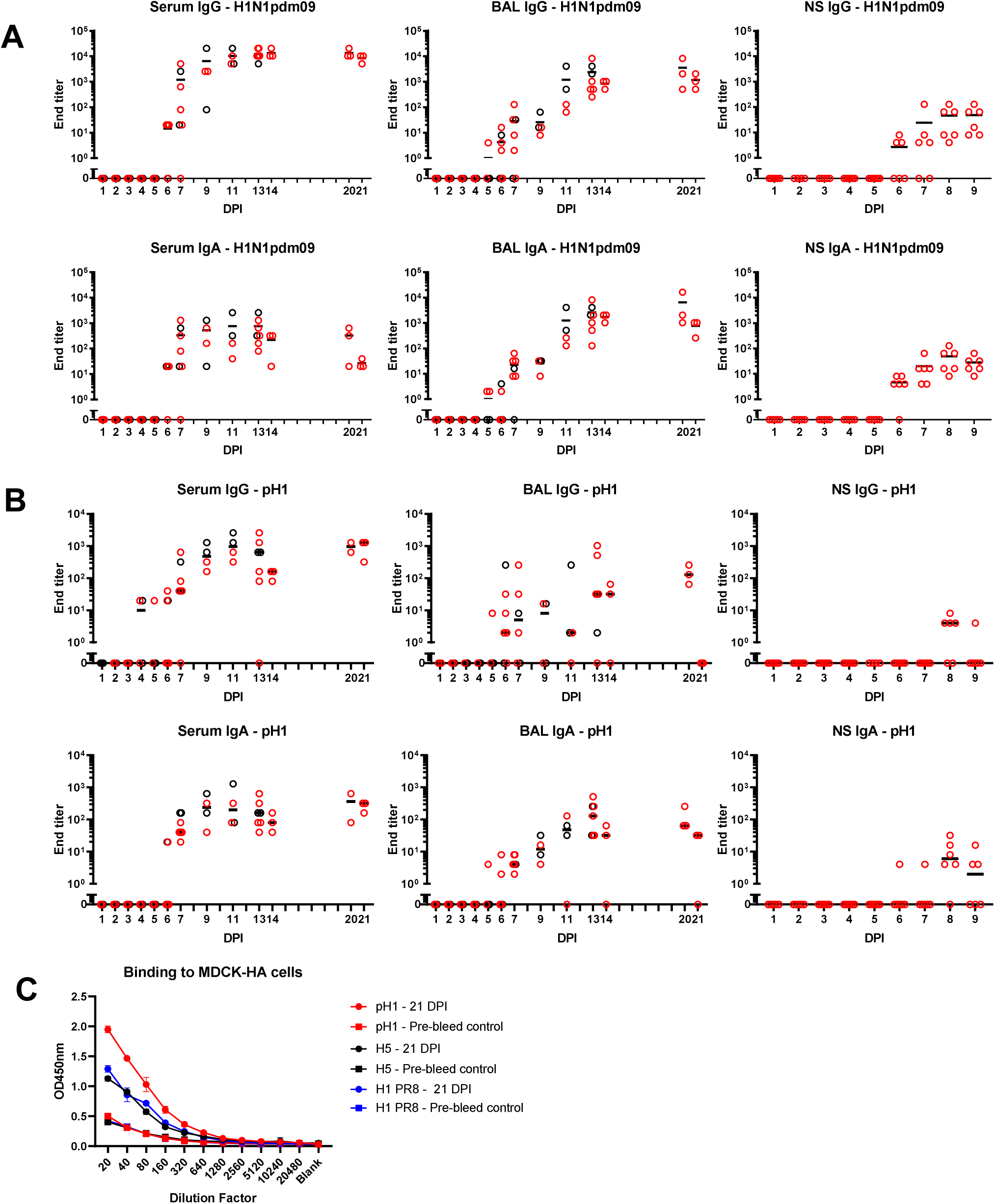
Ab ELISA responses and binding to MDCK-HA expressing cells. Influenza H1N1pdm09 virus specific IgA and IgG **(A)** and haemagglutinin (pHA) specific **(B)** responses in serum, BAL and nasal swabs (NS) were determined by ELISA and shown as black (for OB) and red (for BM) circles. **(C) B**inding of serum at 21 DPI to MDCK-pH1, MDCK-H1 PR8 and MDCK-H5 expressing cells.

To assess the breadth and cross-reactivity of the Ab, we tested the binding of sera from 21 DPI to MDCK cells expressing pH1, H5 (from A/Vietnam/1203/2004) and HA from PR8 in which, unlike in ELISA, the natural conformation of HA is maintained. There was strong binding to the MDCK expressing pH1 and weaker binding to H5 and HA from PR8 suggesting that H1N1pdm09 induces cross reactive responses to other group 1 H1 and H5 viruses (**Fig. 7C)**.

The function of antibodies in serum and BAL was tested by microneutralization (MN) assessing inhibition of virus entry, inhibition of hemagglutination (HAI) and inhibition of neuraminidase activity by enzyme-linked lectin assay (ELLA) (**Fig. 8A**). MN was first detected in serum at 5 or 6 DPI mirroring Ab production in the tissues, increasing to 1:140 at 11 DPI at and 1:480 at 21 DPI. HAI and ELLA followed a similar pattern reaching 1:746 HAI or 1:160 ELLA at 21 DPI. BAL showed much lower MN, HAI and ELLA responses compared to serum. MN and ELLA titres in BAL peaked at 13 DPI and were maintained until 20 DPI. HAI reached a peak at 11 DPI and was undetectable at DPI 21. No significant differences in MN, HAI and ELLA in the upper asymptote, rate of increase in titre or time of maximum increase between OB and BM animals were detected

**Figure 8.**
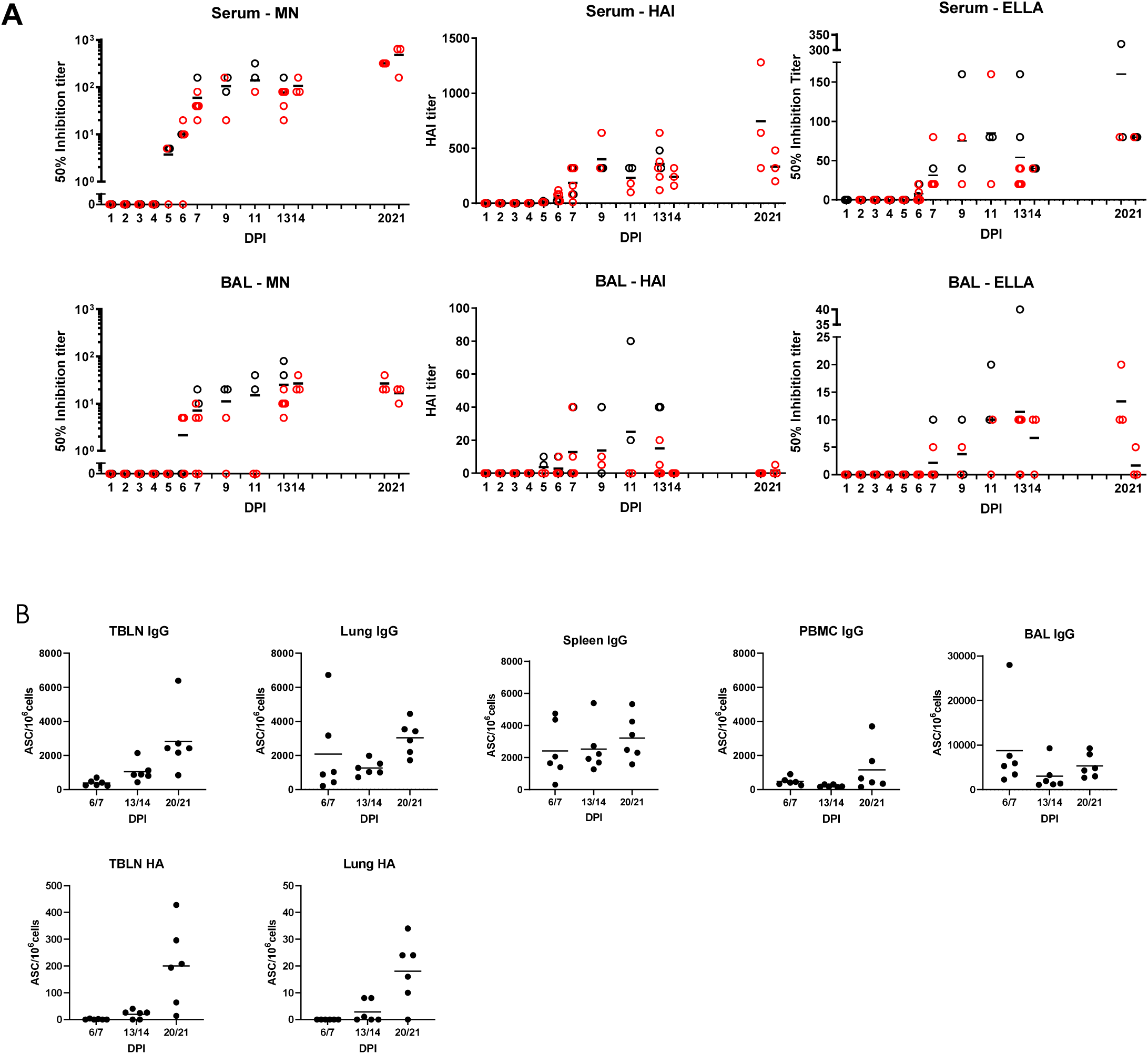
Ab function and antibody secreting cells in tissues. (A) The mean neutralisation (MN), hemagglutination inhibition (HAI) and ELLA titers in serum and BAL over time are shown as red (BM) and black (OB) circles. (B) IgG and pHA specific spot forming cells (SFC) were enumerated in blood of animals from experiment BM3 at the indicated time points in TBLN, lung, spleen, BAL and PBMC.

To determine the major sites of Ab production following H1N1pdm09 infection BAL, lung, TBLN, spleen and PBMC from experiment BM3 were tested for total IgG and HA-specific Ab secreting cells (ASC) (**Fig. 8B**). IgG producing cells were detected in all tissues with a trend for increasing numbers over time up to 21 DPI. TBLN showed the highest frequency of HA specific ASC reaching 200 ASC/10^6^cells at 20/21 DPI. Lung demonstrated a similar pattern but with 18 ASC/10^6^cells at 20/21 DPI.

In summary a strong Ab response was detected in serum, which was dominated by IgG, while in BAL the ELISA titers of IgG and IgA were comparable. Antibodies cross reacted with HA from H1 and H5 viruses. Microneutralization, HAI and ELLA titers were much higher in serum than BAL. HA specific ASC were detected in TBLN and lung. No differences were observed in the Ab responses between OB and BM animals.

## Discussion

In this study we investigated the kinetic and magnitude of T cell and Ab responses in respiratory tissues and blood in outbred Landrace x Hampshire cross and inbred Babraham pigs following H1N1pdm09 infection. The relationship between these parameters and the virus load is illustrated in **Fig. 9**. After experimental infection with H1N1pdm09 virus shedding plateaued between 1 and 4 - 5 DPI, followed by a steep decline so that by 9 DPI no virus could be detected in any animal. An *ex vivo* γδ cell IFNγ and TNF response was apparent from 2 DPI, although this declined by 7 DPI. In contrast, virus specific IFNγ producing γδ cells were detected at 7 DPI and maintained to 13 DPI. Significant virus specific CD4 and CD8 T cell response were present at 6 DPI. Similarly, virus-specific IgG and IgA were detected in serum and BAL at 5 - 6 DPI by which time the viral load had declined by 2-3 logs. By the time of the peak of the T cell and Ab responses (9-14 DPI), no virus was detectable. These kinetics suggest that innate mechanisms, including perhaps early γδ cell cytokine secretion, contain viral replication at a plateau level in the first 4-5 days post infection, while adaptive T and Ab responses contribute to the complete clearance of virus after 5 DPI in primary infection and prevent future infections by a more rapid secondary immune response.

**Figure 9.**
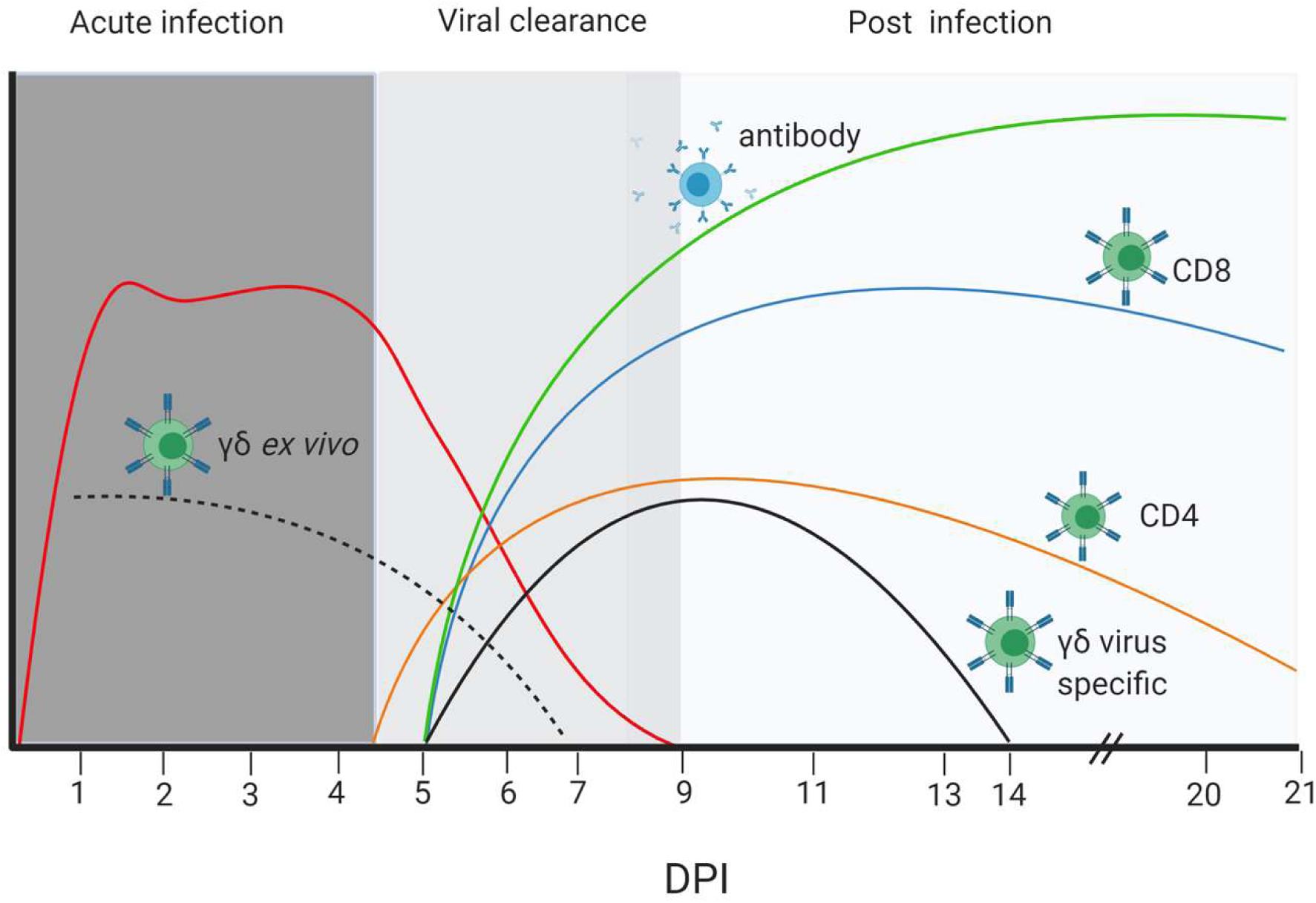
Dynamics of viral load, T cell and Ab responses. Stylised presentation of kinetics and magnitude of viral load and immune responses. The different colour lines represent the antibody and cellular responses as indicated, and the red line the virus load. The dotted black line represents *ex vivo* cytokine production by γδ cells, while the solid black line is the cytokine production by γδ cells re-stimulated with H1N1pdm09 virus *in vitro*. The figure was created with BioRender.com.

Similar kinetics of adaptive T cell responses have been reported in mice, with antigen specific cells detected as early 4-5 days post infection, increasing in number between 5 −12 DPI in lung tissues (Lawrence et al., 2005; Miao et al., 2010). Experiments in mice have shown that depletion of B or CD8 T cells results in delayed clearance of IAV (Eichelberger et al., 1991; Bender et al., 1992; Sarawar et al., 1994; Graham and Braciale, 1997). CD4 T cells also contribute to control of influenza infection, although depletion of this cell subset alone only slightly delayed viral clearance (Topham et al., 1996; Román et al., 2002; Jelley-Gibbs et al., 2005). The strong CD8 and Ab responses detected in the present study suggests that these cell types are also important for viral clearance in pigs. This could be confirmed by depletions studies or cell transfer in inbred Babraham pigs.

Few studies have analysed in depth the conventional T cell response in pigs. The most comprehensive study showed a low frequency of virus specific IFNγ producing CD4 and CD8 in the lung as early as 4 DPI after H1N2 intratracheal challenge, reaching a peak at 9 DPI, with the highest response in lung compared to TBLN or PBMC (Talker et al., 2016). Here, for the first time, we have analysed the cytokine responses in BAL as well as lung interstitial tissues, which showed a similar kinetic. However, the response in the BAL was much stronger in terms of frequency of cytokine producing T cells. The BAL T cells produced multiple cytokines and more per cell, indicating that they may be most efficient in clearing the virus. Cytokine production differed between CD8 and CD4 cells and between BAL, lung and TBLN, perhaps reflecting the extensive tissue compartmentalization in the respiratory tract and differential localization of CD4 and CD8 T cells (Topham and Reilly, 2018). Whether specialized CD4 and CD8 cells are compartmentalized due to the migration of different subsets to specific sites, as has been proposed in mice, or because tissue environments alter cytokine production remains to be established (Strutt et al., 2013).

An important difference between the present study and Talker *et al* is that they used the more pathogenic swine H1N2 virus, which was delivered in a large volume and high dose (15 ml of 10^7^ TCID_50_/ml) intratracheally (Talker et al., 2016). This might explain the stronger and more prolonged lung, TBLN and PBMC responses they observed. The pigs in the present study were infected intranasally with a MAD and the response here was similar to the in-contact animals, suggesting that this method is more similar to natural infection. Furthermore our scintigraphy study also indicates that this method of challenge targets both the upper and lower respiratory tract (Martini, 2020).

We detected a 27 times lower proportion of CD8 antigen-specific cells in the blood compared to BAL. Similarly, antigen-specific CD8 cells responses were much higher in the BAL of patients with H1N1pdm09 compared to blood (Zhao et al., 2012). This indicates that sampling blood is not reflective of the true response in the lung and local tissues, which has implications for the design and analysis of clinical trials for T-cell targeted vaccines. In contrast, CD4 responses were more similar in magnitude in blood and BAL, although less long lived than CD8.

The contributions of γδ cells to lung homeostasis and influenza immunity remain incompletely explored. In pigs, γδ cells comprise up to 50% of lymphocytes in the blood (particularly in young animals) in contrast to humans where they usually represent 1-5% of lymphocytes (Roden et al., 2008; Schwaiger et al., 2019). γδ T cell have previously been reported to increase late after IAV infection in mice, although an early increase in γδ cells in mice and pigs has also been reported (Carding et al., 1990; Khatri et al., 2010; Palomino-Segura et al., 2020). Human γδ cells can expand in a TCR-independent manner in response to IAV, and the human Vγ9Vδ2 T cell subset kills IAV-infected A549 airway cells (Li et al., 2013b). Although we did not observe a significant increase in γδ cells after H1N1pdm09 infection, we showed that γδ cells produce IFNγ and TNF as early as day 2 post infection *ex vivo*, in agreement with studies in mice (Xue et al., 2017). γδ cells are a major source for IL-17 production, which has been shown to play a role in IAV infection, but we detected only low levels of IL-17A after H1N1pdm09 stimulation of BAL cells (Crowe et al., 2009; Li et al., 2012; Palomino-Segura et al., 2020). Surprisingly, we demonstrated that *in vitro* stimulation with H1N1pdm09 induces IFNγ and TNF production in CD2^+^ γδ T cells from 7 to 13 DPI. This is reminiscent of an adaptive T cell response. Recombinant hemagglutinin from H5N1 has been previously demonstrated to activate human PBMC γδ T cells *in vitro* and this was not mediated by TCR or pattern recognition receptors (Lu et al., 2013). Further studies will elucidate the mechanisms of cytokine induction and whether it is TCR dependent.

H1N1pdm09 infection was characterised by high IgG and IgA titers in serum and BAL, and a detectable antibody titer in nasal swabs. The IgG titer was higher than IgA in serum, while similar levels of IgA and IgG were detected in BAL and nasal swabs, suggesting local production of this isotype or more efficient translocation. Neutralization, HAI and neuraminidase inhibition titers peaked at 11 - 21 DPI. Our findings are in agreement with previous studies showing that in experimentally H1N1 infected pigs HA-specific antibodies peaked at 2-3 weeks (Larsen et al., 2000). Similarly we detected HA-specific antibody-secreting cells in the local TBLN and lung tissues, but not PBMC (Larsen et al., 2000). However, it might be that antibody-secreting cells are largely lost in these liquid nitrogen frozen and thawed samples.

Despite centuries of agricultural selective breeding, the pig has maintained a significant level of SLA genetic diversity, with 227 class I and 211 class II alleles identified for *Sus scrofa* in the Immuno‐Polymorphism Database (IPD) MHC database to date, making analysis of the fine specificity of immune responses extremely difficult (Maccari et al., 2017). The inbred Babraham line of pigs, on the other hand, is SLA homozygous for class I SLA‐ 1*14:02; SLA‐2*11:04 and SLA‐3*04:03 and class II DRB1-*05:01, DQA-*01:03 and DQB1-*08:01 (Schwartz et al., 2018). This homozygosity enabled the use of peptide-SLA tetramers to the dominant NP antigen to track the CD8 response in tissues in this study (Tungatt et al., 2018). The αβ, γδ T-cell and Ab responses in the OB and BM animals were comparable, although there was lower proportion of CD8 cells and higher proportion of γδ cells in the BM pigs. This may be due to a genetic difference, although it may also be a result of different housing conditions, since the Babraham pigs are maintained under specific pathogen free conditions, whereas the outbred pigs were obtained from a commercial breeder.

Our detailed analysis of immune responses in pigs showed that the viral load is contained in the period before the adaptive response is detectable, indicating the importance of innate immune mechanisms in influenza infection. As in other species however it appears that the adaptive response is essential for elimination of virus. BAL contains the most highly activated CD8, CD4 and γδ cells producing large amounts of cytokines, which likely contribute to clearance of virus. We further show clear differences between the function of CD4, CD8 and γδ T cells between the lung, BAL and TBLN, while the blood is a poor representation of the local immune response. The immune response in the Babraham pig following H1N1pdm09 influenza infection was comparable to that of outbred animals. The availability of fine grain immunologic tools in Babraham pigs will allow the unraveling of immune mechanisms and confirm and extend findings in outbred populations.

## Acknowledgements

We are grateful to animal staff at Langford School of Veterinary for providing excellent animal care. We are grateful to Alain Townsend for providing recombinant HAs and H7N1 virus. We thank APHA for providing the swine A/Sw/Eng/1353/09 influenza virus strain (DEFRA surveillance programme SW3401).

## Authors contribution

BC, MB, ET conceived, designed and coordinated the study. BC, MB, ET, ME, AM, EP, EV, BP, VM OF, RH, AT, RB, SM designed and performed experiments, processed samples and analyzed the data. SG performed statistical analysis. AF and AS generated SLA tetramers. ET, ME, EV, AM wrote and revised the manuscript and figures. All authors reviewed the manuscript.

---

**Supplementary Figure 1.**
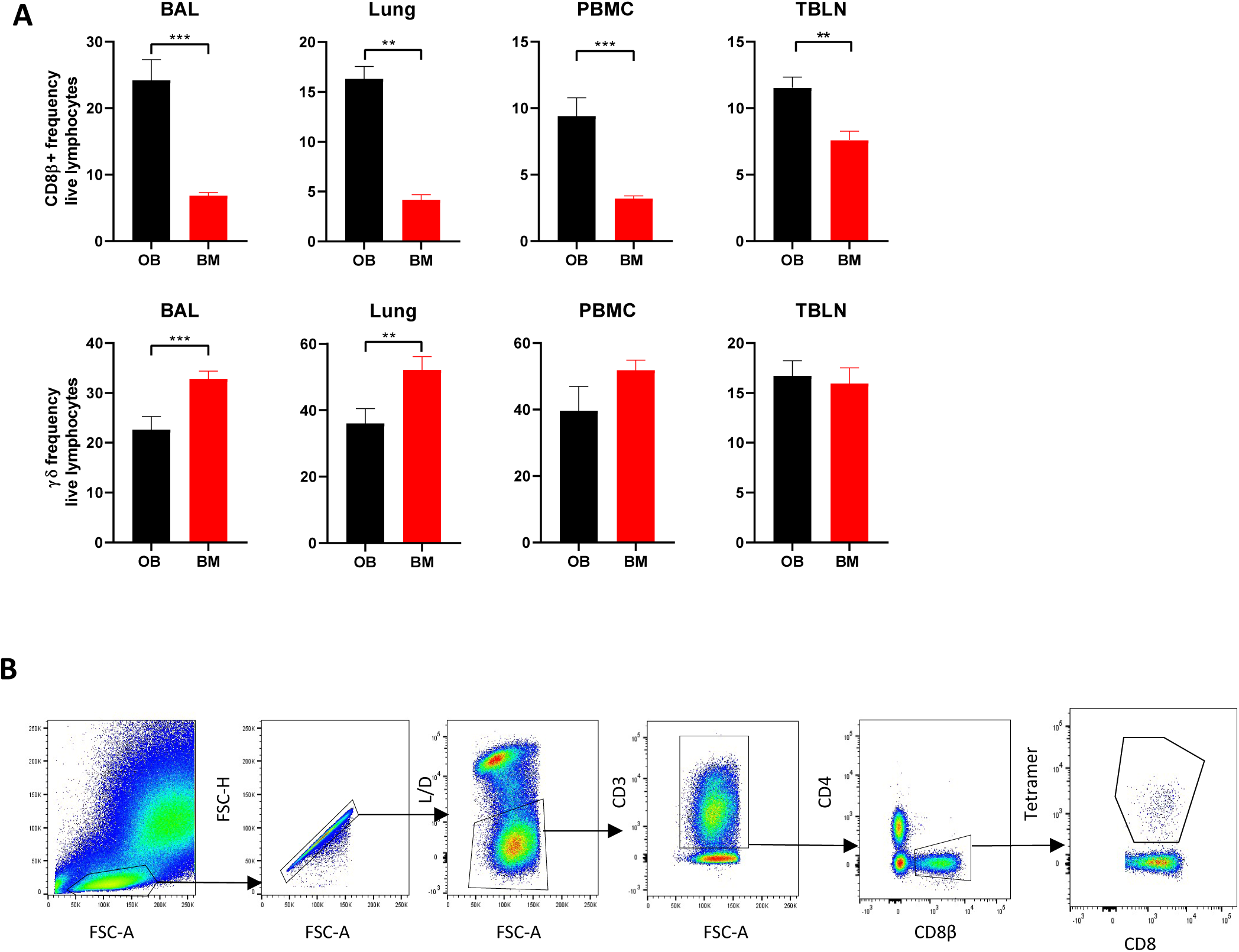
Proportion of CD8 cells in OB and BM and gating strategy for tetramer identification. (**A)** Comparison of proportions of CD8β and γδ cells in control uninfected outbred (experimentsOB1 and OB2) and inbred Babraham (experiments BM1 and BM2) pigs. Asterisks denote * p≤ 0.05, ** ≤ 0.005, *** ≤ 0.0001. (**B)** Gating strategy for identification of NP_290-298_ tetramer T cells. BAL, lung, TBLN and PBMC were stained with the relevant antibodies. Lymphocytes were gated by light scatter, followed by exclusion of doublets and dead cells. CD3+, CD4-CD8β+ T cells were gates and the Tetramer+ population enumerated as a percentage of CD8β+ T cells.

**Supplementary Figure 2:**
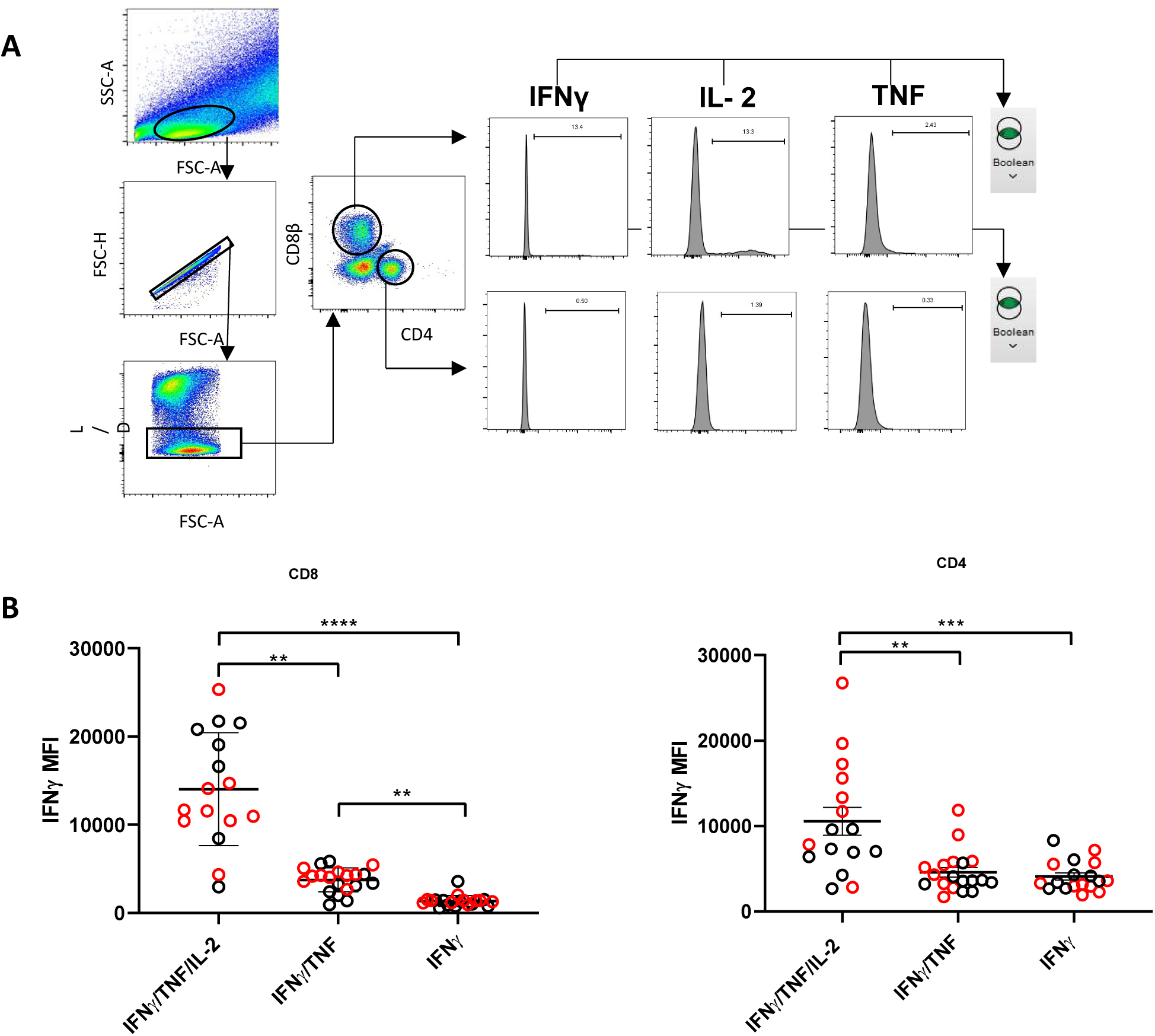
Gating strategy and cytokine production by CD8 and CD4 cells. **(A)** Gating strategy for identification of CD8β and CD4 cytokine producing cells. BAL, lung, TBLN and PBMC were stimulated with H1N1pdm09, followed by intracytoplasmic staining. Lymphocytes were gated by light scatter and further sub-gated for exclusion of dead cells with a live/ dead discrimination dye. CD8β or CD4 cells were gated and expression of IFNγ, TNF or IL-2 was determined by histogram. Boolean gating of all cytokine positive cells was performed and cytokine responses determined by summing the total cytokine production in single, double and triple producing cells. **(B)** MFI of IFNγ secretion was determined in CD8β and CD4 cells identified as IFNγ single, IFNγ/TNF double or IFNγ/TNF/IL-2 triple secreting T cells. IFNγ MFI is plotted for BM (red circles) and OB (black circles) animal. Asterisk indicates *p≤0.05, **p≤0.01 and ***p ≤ 0.001.

**Supplementary Figure 3.**
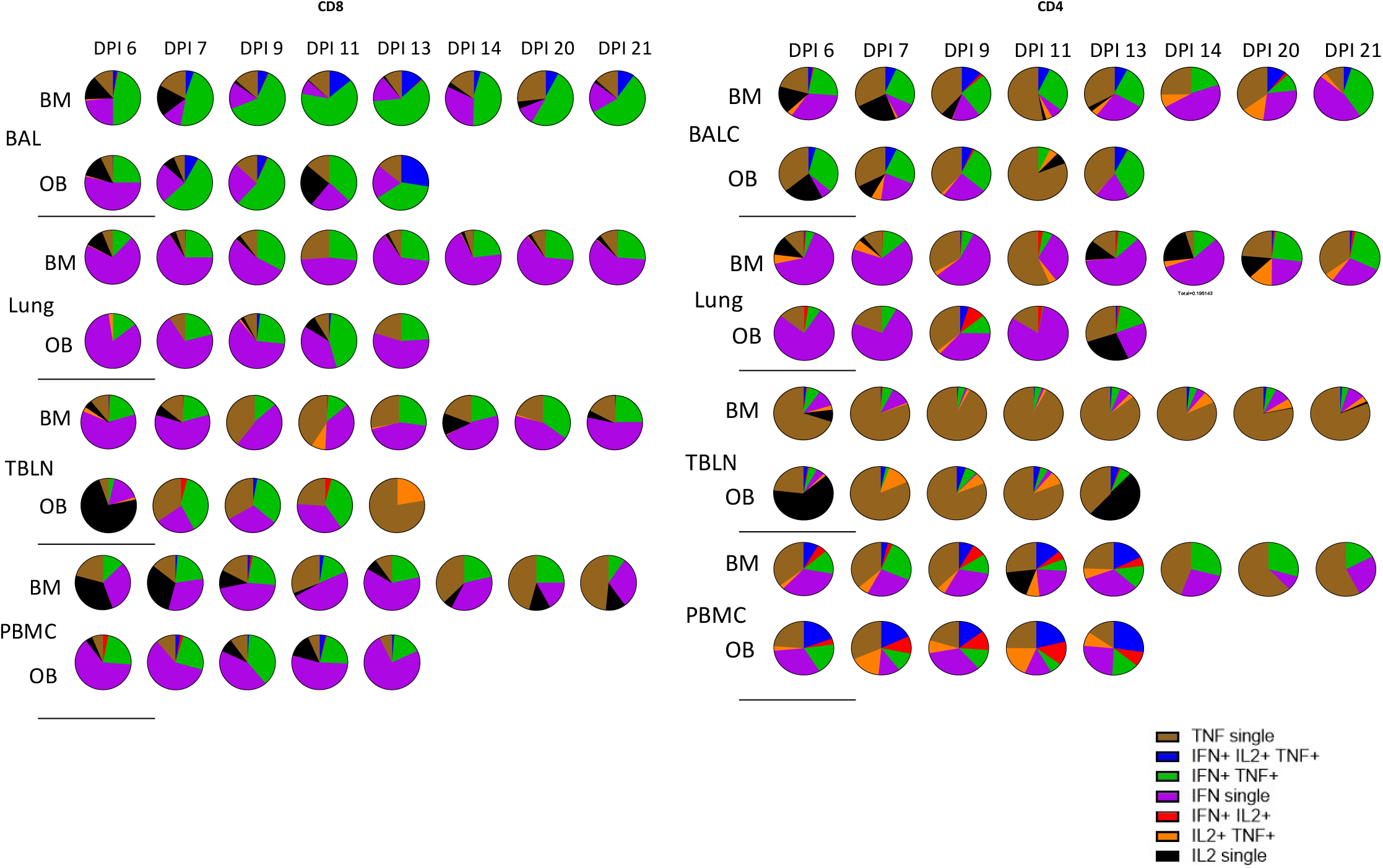
Cytokine production by CD8β and CD4 cells. Boolean gating of cytokine producing CD8 and CD4 T cells in tissues. Pie charts depict the proportion of single, double and triple IFNγ, IL-2 and TNF cytokine producing cells as a proportion of the total cytokine production in BAL, lung, TBLN and PBMC. Cytokine response for 6, 7, 9, 13 and 21 DPI for both OB and BM are shown. Medium control was subtracted from each stimulated population.

**Supplementary Figure 4:**
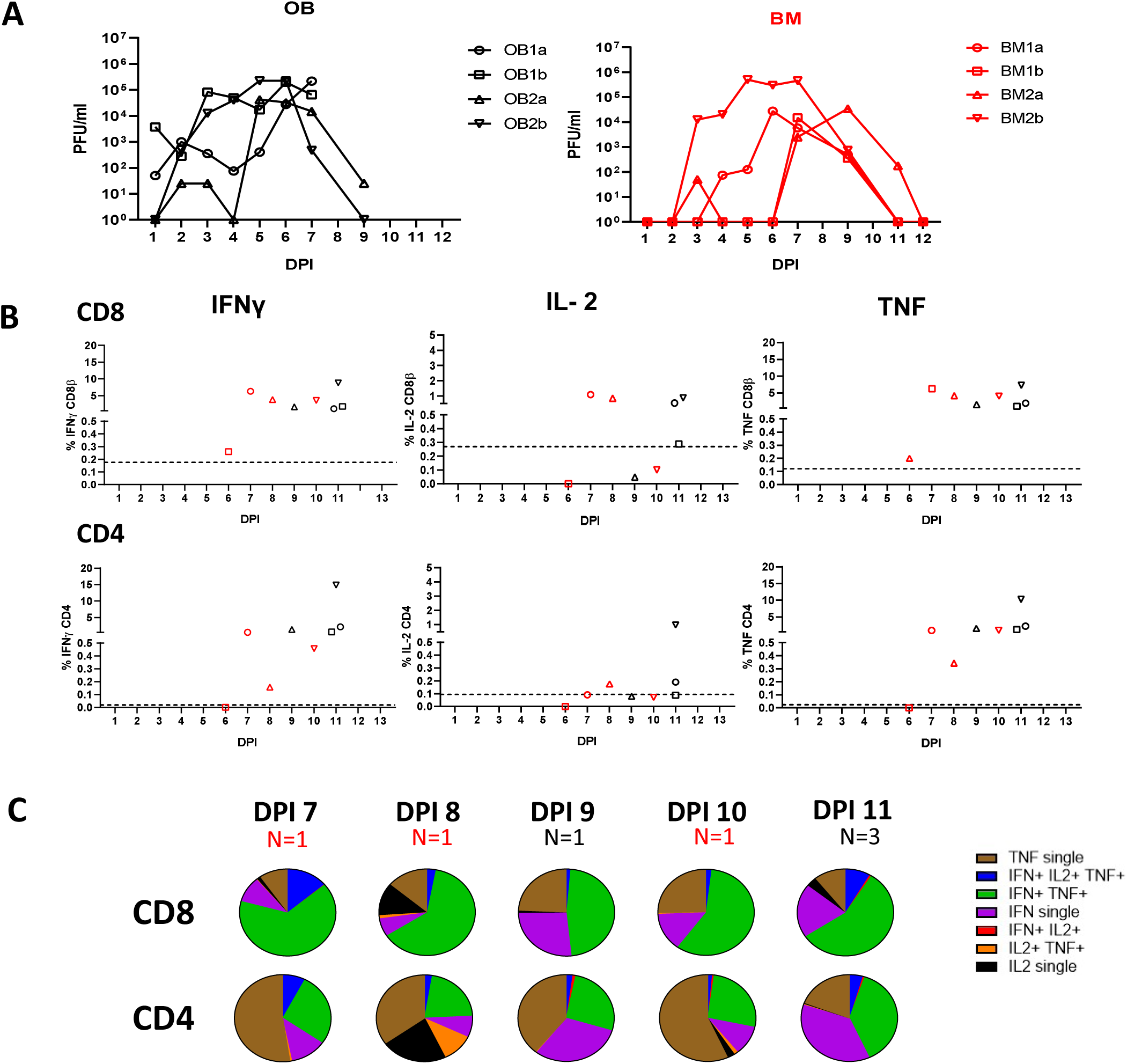
CD8β and CD4 cytokine response in BAL of in-contact animals. **(A)** Virus load in in-contact animals was determined by plaque assay of daily nasal swabs at the indicated time points for outbred (black) and inbred Babraham (red) pigs. **(B)** BAL CD4 and CD8 cytokine responses were determined by intracytoplasmic staining and shown as the days post infection counting from the first day the animal shed virus. **(C)** Pie charts show the proportion of single, double and triple cytokine secreting CD8 T cells for IFNγ, TNF and IL-2.

**Supplementary Figure 5.**
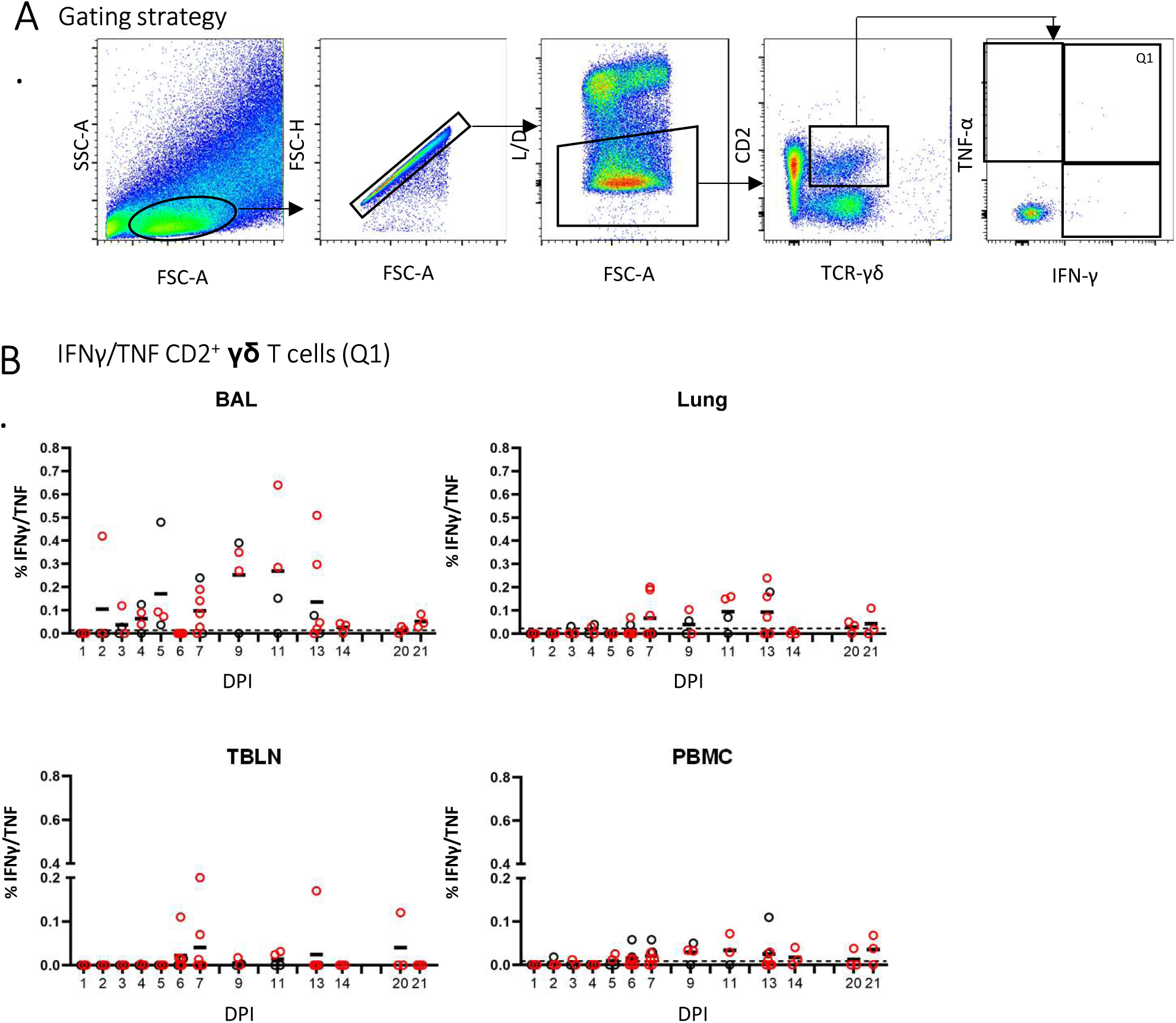
Double cytokine producing γδ cells after H1N1pdmd09 stimulation. **(A)** Lymphocytes were gated by light scatter properties and further sub-gated for exclusion of doublets and dead cells with a live/ dead discrimination dye. The CD2^+^ γδ T cells were analyzed for co-production of IFNγ/TNF (Q1). **(B)** Frequency of IFNγ/TNF co-producing CD2^+^ γδ T cells in outbred (black circles) and inbred (red circles) pigs following influenza infection. BAL, lung, TBLN and PBMC cells were stimulated overnight with H1N1pdm09 followed by intracellular cytokine staining. DPI 1 to 7, 9, 11 and 13 each show results from 2 outbred and 2 inbred pigs. DPI 6, 7, 13, 14, 20 and 21 also include results from 3 additional inbred pigs. The mean of the 22 uninfected control animals is represented by a dotted line.

**Supplementary Figure 4:**
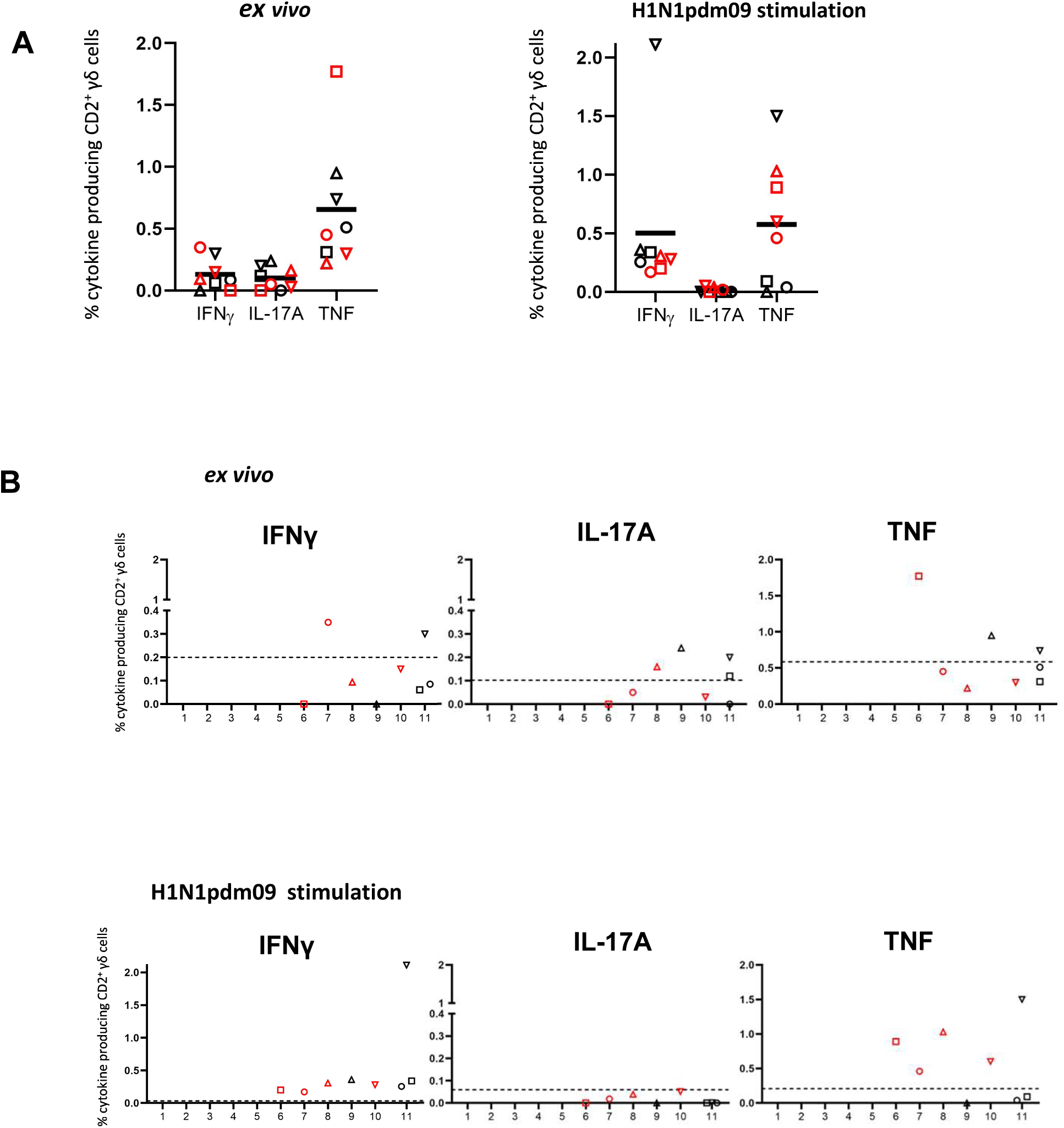
BAL cytokine response in γδ cells of in-contact animals. **(A)** Proportions of IFNγ, TNF and IL-17 cytokine producing cells in all in-contacts and **(B)** at each time point after infection for the individual in-contact animals shown as the days post infection counting from the first day the animal shed virus.

